# Phosphodiesterase 3A modulators sensitize tumor cells to Bcl-xL and Bcl-2/Bcl-xL inhibitors

**DOI:** 10.1101/2025.04.10.648087

**Authors:** Kirsi Toivanen, Astrid Murumägi, Agnieszka Wozniak, Karo Wyns, Chao-Chi Wang, Luna De Sutter, Mariliina Arjama, Nanna Merikoski, Sami Salmikangas, Omar Youssef, Jorma Isola, Olli Kallioniemi, Patrick Schöffski, Tom Böhling, Harri Sihto

## Abstract

Phosphodiesterase 3A (PDE3A) modulators are emerging anticancer agents for tumors that co-express PDE3A and Schlafen 12 (SLFN12), promoting the formation of a cytotoxic PDE3A-SLFN12 complex. The effect correlates with PDE3A and SLFN12 expression levels. In this study, we conducted a high-throughput screening of 526 compounds in three sarcoma cell lines to identify synergistic or antagonistic interactions with PDE3A modulators. Synergistic mechanisms were further investigated in cell lines and in patient-derived xenograft models of leiomyosarcoma and myxofibrosarcoma in mice. Several drug classes showed potential synergy, and PDE3A modulators sensitized cells to otherwise ineffective Bcl-2/xL inhibitors. The combination of a PDE3A modulator and a Bcl-xL inhibitor induced tumor regression in a patient-derived leiomyosarcoma model *in vivo*. Our findings reveal that PDE3A modulators synergized with several inhibitors of key cancer pathways, offering promising potential for future treatments. With further research, PDE3A modulators and combination therapies could provide effective targeted treatment options for tumors expressing both PDE3A and SLFN12, paving the way for innovative and hopeful advancements in cancer treatment.

## Introduction

Phosphodiesterase 3A (PDE3A) modulators are structurally diverse compounds that exhibit a unique molecular glue capability [1-6]. Also referred to as velcrins [5, 7, 8], these compounds can form a complex between two proteins, PDE3A and Schlafen 12 (SLFN12), inducing toxicity in cells that express both proteins [1-5, 9]. The cytotoxic effect is dose-dependent [2-4, 10-16], and correlates with both PDE3A and SLFN12 expression levels [2, 7, 15-17]. PDE3A modulators reduce cell viability across multiple cancer cell lines, including gastrointestinal stromal tumor (GIST) [14], liposarcoma [9], embryonic rhabdomyosarcoma [11], alveolar soft part sarcoma [18], hepatocellular carcinoma [17, 19], various adenocarcinomas [1-4, 6, 8, 10-13, 15-17], neuroglioma [4, 6, 17], glioblastoma [7, 15], primitive neuroectodermal tumor [16], melanoma [15, 17], and erythroleukemia [15]. PDE3A modulators have also been reported to have antitumor effects *in vivo* in mice inoculated with cell lines of GIST [14], lung adenocarcinoma [16], cervical adenocarcinoma [1, 4], hepatocellular carcinoma [19], primitive neuroectodermal tumor [16], neuroglioma [4, 17], and glioblastoma [7], as well as in patient-derived xenografts (PDX) of GIST [14, 16], sarcoma [16], ovarian cancer [16], lung cancer [16], glioblastoma [7], and melanoma [16], further supporting their clinical potential.

Despite their structural diversity, PDE3A modulators induce a similar butterfly-shaped heterotetramer of PDE3A and SLFN12 [1, 5]. These two cytosolic proteins, which otherwise do not typically interact with each other, are brought together by the PDE3A modulator [5, 15, 17]. The modulators first attach to the catalytic sites of a PDE3A dimer, subsequently binding to a dimer of SLFN12 [5, 15], with multiple hydrogen bonds and van der Waals interactions holding the macromolecule together [1, 5]. SLFN12, which is otherwise rapidly ubiquitinated and degraded by the proteasome, is stabilized [3, 5, 16] and dephosphorylated [6], increasing its ability to degrade rRNA and tRNA [5, 8]. This ultimately results in decreased protein translation and increased endoplasmic reticulum stress [3, 5, 8, 16]. Mutations at the complex-forming sites of these two proteins have been shown to reduce sensitivity to PDE3A modulators [4, 15].

The need for precision medicine approaches is particularly evident in sarcomas, as standard treatment involves surgical resection, often accompanied by radiation or chemotherapy, with outcomes largely dependent on tumor subtype. PDE3A modulators could serve as precision medicine for sarcomas that eminently express PDE3A, specifically GIST, leiomyosarcoma, and myxoid liposarcoma [9, 12-14]. However, in cases where the effect results in cell cycle arrest rather than apoptosis due to low or moderate PDE3A and SLFN12 expression levels [17], combining PDE3A modulators with synergistic drugs may enhance their therapeutic potential, even when they are less effective as single agents.

In this study, we investigated 527 drug combinations with two PDE3A modulators to evaluate their synergistic effects in three PDE3A- and SLFN12-positive sarcoma cell lines: one GIST cell line and two liposarcoma cell lines. This study aimed to: i) identify drug partners that could enhance PDE3A modulator efficacy; ii) examine the molecular pathways contributing to synergistic interactions; and iii) evaluate synergistic effects *in vivo* using PDX sarcoma models.

## Methods

The following cell lines, GOT3 (RRID: CVCL_M819), and MLS402-91 (RRID: CVCL_S813) were kindly provided by Prof. Pierre Åman (University of Gothenburg, Gothenburg, Sweden), GIST48 (RRID: CVCL_7041) and GIST882 (RRID: CVCL_7044) by Dr. Jonathan Fletcher (Harvard Medical School, Boston, MA, USA); and the SA4 (RRID: CVCL_8910) cell line by Dr. Kjetil Boye (Oslo University Hospital, Oslo, Norway). GIST-T1 cell line (RRID: CVCL_4976, Cat. No. PMC-GIST01C) was purchased from Cosmo Bio (Tokyo, Japan), and A2058 (RRID: CVCL_1059, Cat. No. CRL-3601), H4 (RRID: CVCL_1239, Cat. No. HTB-148), and CHL-1 (RRID: CVCL_1122, Cat. No. CRL-9446) cell lines were purchased from ATCC (Manassas, VA, USA). Sequencing verified known *KIT* mutations; exon 13 p.K642E in GIST882, and *KIT* exon 11 p.V560D and exon 17 p.D820A in GIST48. HPV E7 protein was not detected from SA4 cell line by western blot, indicating no HeLa contamination. All cell lines were authenticated using short-tandem-repeat (STR) profiling in the genotyping lab of the Institute for Molecular Medicine Finland (FIMM) Technology Centre (Helsinki, Finland), tested negative for *Mycoplasma*, and cultured in a humidified, 5% CO2 atmosphere at +37 °C.

GIST-T1, A2058, H4, and CHL-1 cell lines were cultured in Dulbecco’s modified Eagle’s medium (Cat. No. 21969-035, Gibco, Thermo Fisher Scientific, Waltham, MA, USA) with 10% inactivated fetal bovine serum (iFBS, Cat. No. 10270-106, Gibco), 100 U/mL penicillin, 100 U/mL streptomycin, and 0.03 mg/mL L-glutamine (Pen Strep Glut, Cat. No. 10378-016, Gibco). Other cell lines were cultured in RPMI Medium 1640 (Cat. No. 11530586, Gibco) with Pen Strep Glut (Gibco) and 5% (MLS402-9), 10% (SA4 and GOT3), or 20% (GIST48, and GIST882) iFBS (Gibco).

### High-throughput compound screening

High-throughput drug sensitivity and resistance tests (DSRT) with 526 FDA-approved or investigational compounds and probes were performed in FIMM as previously described [20]. DSRT was performed for screening synergy combinations in GIST882, SA4, and GOT3 cells with and without PDE3A modulators, 100 nM anagrelide (ANA) and 40 nM BAY 2666605 (BAY). The experiments included one triple-drug combination of cytarabine, idarubicin, and PDE3A modulator, resulting in 527 total combinations. Briefly, dissolved compounds (in DMSO or water, depending on solubility) were dispensed covering a 10,000-fold concentration range into Corning 384-well plates (Cat. No. CLS3573, Merck KGaA, Darmstadt, Germany) that were then stored in pressurized StoragePods® (Roylan Developments Ltd., Surrey, UK). Plated cells (1,000 cells/well) were incubated with the compounds for 72 hours at +37 °C, prior to measuring cell viabilities with CellTiter-Glo® (Cat. No. G7572, Promega, Madison, WI, USA) and microplate reader PHERAstar (BMG LABTECH, Ortenberg, Germany). Drug responses were evaluated using the area under the dose-response curve, quantified as Drug Sensitivity Scores (DSS) [21], which were calculated using the data analysis tool Breeze (FIMM) [22]. Synergistic and antagonistic effects were measured for each compound in each cell line using ΔDSS values, obtained by subtracting the DSS value of the compound from the corresponding DSS value of the combination; combinations that increased the DSS value compared to the drug alone were considered potentially synergistic, and vice versa. Synergy and antagonism were further assessed among the cell lines by selecting compounds whose ΔDSS values coincided with positive or negative results in at least two cell lines per PDE3A modulator.

Based on the ΔDSS results, 10 compounds were selected for combination testing with ANA in a 7×7 concentration matrix: A-1155463, A-1331852, navitoclax, ridaforolimus, everolimus, sirolimus, temsirolimus, PF-03758309, quisinostat, and pevonedistat. Synergy was assessed using the zero interaction potency model (ZIP score) calculated with SynergyFinder (FIMM) [23]. Highly positive ZIP scores indicate synergy, whereas near zero values indicate an additive effect and negative values antagonism.

### Cell viability

The following drugs were used in drug efficacy experiments: anagrelide hydrochloride (CAS No. 58579-51-4, Cat No. 735554, Lancrix, Shanghai, China), cilostazol (CAS No. 73963-72-1, Cat. No. C0737, Sigma-Aldrich, Saint Louis, MO, USA), A-1155463 (CAS No. 1235034-55-5, Cat. No. HY-19725, MedChemExpress, Monmouth Junction, NJ, USA), A-1331852 (CAS No. 1430844-80-6, Cat. No. HY-19741, MedChemExpress), navitoclax (CAS No. 923564-51-6, Cat. No. S1001, Selleckchem, Houston, TX, USA), 6-(4-(diethylamino)-3-nitrophenyl)-5-methyl-4,5-dihydropyridazin-3(2H)-one (DNMDP; CAS No. 328104-79-6, Cat. No. HY-W028690, MedChemExpress), and BAY 2666605 (CAS No. 2275774-60-0, Cat. No. HY-145924, MedChemExpress). All drugs were dissolved in DMSO and further diluted with PBS. Corresponding DMSO concentration in cell media was used as control treatment. Based on the observed synergy concentration distributions (Supplementary Figures S1–S3), 100 nM concentrations for ANA, A-1155463, and navitoclax were selected for subsequent cell viability experiments. A 10 nM concentration was chosen for DNMDP as it approximately corresponded the 100 nM ANA concentration (Supplementary Figure S4).

Cells were seeded on ViewPlate-96, White 96-well Microplate with Clear Bottom (Cat. No. 6005181, Revvity, Waltham, MA, USA) 24 hours before replacing the medium with drug-containing medium. Varying cell seeding amounts were used to achieve 70–80% confluence in the control treatment on the final measurement day. Eight replicate wells were used for each treatment and cell line. Cell viability was measured using CellTiter-Glo® (Promega), and luminescence was recorded with Hidex Sense microplate reader (Hidex Oy, Turku, Finland). The synergy results presented were obtained from two independent experiments. For sequential treatment, cells were treated with first drug for 48 hours, after which media with second drug was changed for 48 hours.

### Western blotting

Cells were lysed with M-PER™ Mammalian Protein Extraction Reagent (Cat. No. 78501, Thermo Fisher Scientific) containing HALT™ protease (Cat. No. 78429, Thermo Fisher Scientific) and phosphatase (Cat. No. 78420, Thermo Fisher Scientific) inhibitor cocktails, following the manufacturer’s instructions. Protein detection with western blot was performed as previously described [9]. The following antibodies were used; polyclonal rabbit anti-PDE3A (1:1,000, Cat. No. HPA014492, Sigma-Aldrich), monoclonal rabbit anti-SLFN12 (1:500, Cat. No. ab234418, Abcam, Cambridge, UK), polyclonal rabbit anti-PARP (1:1,000, Cat. No. 9542S, Cell Signaling Technology, Beverly, MA, USA), polyclonal rabbit anti-caspase-3 (1:1,000, Cat. No. 9662, Cell Signaling Technology), polyclonal rabbit anti-Mcl-1 (1:1,000, Cat. No. 4572, Cell Signaling Technology), monoclonal rabbit anti-Bcl-xL (clone 54H6, 1:1,000, Cat. No. 2764, Cell Signaling Technology), polyclonal rabbit anti-Bcl-2 (1:1,000, Cat. No. 59348, Abcam), and polyclonal rabbit anti-cytoskeletal actin (1:150,000, Cat. No. A300-491A, Bethyl Laboratories, Montgomery, TX, USA). Band densitometry was measured using ImageJ image analysis software and normalized to cytoskeletal actin expression.

### SLFN12 siRNA

Forward transfection was performed 24 hours after cells were seeded using two SLFN12 siRNA products; Hs_SLFN12_1 Flexitube siRNA (NM_018042, 2530 bp, GeneGlobe ID - SI04132233|S0, Cat. No. 1027415, QIAGEN, Venlo, The Netherlands), and Hs_SLFN12_2 Flexitube siRNA (NM_018042, 2530 bp, GeneGlobe ID - SI04160604|S0, Cat. No. 1027415, QIAGEN). First, siRNA-RNAiMAX complexes were generated by combining the freshly prepared siRNA solutions (0.2 µM of siRNA in Opti-MEM™ Reduced Serum Medium (Cat. No. 31985070, Gibco)) and RNAiMax dilution (5% (v/v) Lipofectamine RNAiMAX Transfection Reagent (Cat. No. 13778150, Invitrogen, Thermo Fisher Scientific) in Opti-MEM™ medium) in a 1:1 ratio. With SA4 cells, the used RNAiMAX concentration was 10% (v/v). After a 5–10 min incubation, siRNA-RNAiMAX complexes were added 10 µL/well to medium. Cells were exposed to siRNA products for 24 hours, after which cell mediums were replaced to mediums containing 0.001% DMSO, 100 nM ANA, 100 nM A-1155463, or both of the compounds.

Cell viabilities measured with CellTiter-Glo® (Promega) on the following day after siRNA treatment were considered as baseline (day 0), to which viabilities measured on the following days were compared, indicating a relative change in cell viability percentage. RNAiMAX-treated cells and a scrambled siRNA product were used as controls (ON-TARGETplus™ Control Pool Non-targeting pool, Cat. No. D-001810-10-05, Dharmacon™, Lafayette, CO, USA).

### TaqMan Array Human Apoptosis

After cells were treated with 0.001% DMSO or 100 nM ANA for 24 hours, RNA was extracted using NucleoSpin RNA (Cat. No. 740955.50, Macherey-Nagel, Düren, Germany) following the manufacturer’s instructions. RNA samples were eluted in 20 µL of nuclease-free water, and after measuring RNA concentration, they were diluted to a concentration of 100 ng/µL. cDNA was then synthesized from RNA using SuperScript VILO cDNA Synthesis Kit (Cat. No. 11754250, invitrogen, Thermo Fisher Scientific), according to manufacturer’s instructions, in 20 µL reaction volumes.

Real-time qPCR was performed in 20 µL reaction volumes on TaqMan Array Human Apoptosis assay plates (Cat. No. 4414072, Thermo Fisher Scientific) with FastStart TaqMan Probe Master Mix (Cat. No. 04673417001, Roche, Basel, Switzerland) using the CFX96 Touch Real-Time PCR detection system (Bio-Rad, Hercules, CA, USA), under the following conditions: +95 °C for 10 minutes, followed by 40 cycles of +95 °C for 15□seconds and +60 °C for 1 minute with plate reads during amplification.

Ct values were obtained using CFX Manager Software (Bio-Rad). Due to excessive evaporation, the results from the corner wells of the plate were excluded from the analysis. Among the four housekeeping genes included in the assay, GAPDH was used as the reference gene for normalizing gene expressions. To analyze changes in gene expression, fold change values were computed using the 2^−ΔΔCq^ method for 89 target genes.

### Ethics statement

Human tumor collection and PDX generation was performed at KU Leuven as previously described [24]. Patients provided informed consent for xenografting of their tumor samples, and the establishment of the mouse models, with names constructed in a pseudonymized manner, was approved by the Medical Ethics Committee of the University Hospitals Leuven (project number S53483). The *in vivo* experiments were approved by the Ethics Committee for Laboratory Animal Research, KU Leuven (project number S67961), and conducted in compliance with Belgian/European Union regulations. All the procedures related to animal handling, care, and the treatment in this study were performed in accordance with the ARRIVE guideline [25].

### PDX sarcoma model selection

To screen sarcoma PDX models that could potentially respond to synergy therapy, PDE3A and SLFN12 expression levels were evaluated by immunohistochemistry (IHC) on 5 formalin-fixed paraffin embedded (FFPE) tissue microarrays (TMAs), containing 592 tissue cores from 73 soft-tissue sarcoma PDX models [26]. All IHC staining conditions are given in Supplementary Table S1. PDE3A and SLFN12 staining intensities were scored as 0 (no staining in tumor cells), 1 (weak), 2 (moderate) or 3 (strong). Average IHC staining scores were calculated for each model, derived from the 2–18 cores included in the TMAs. Models were then classified based on these scores as double positive (PDE3A ≥1 and SLFN12 ≥0.25), or negative (PDE3A < 1 or SLFN12 < 0.25), indicating their potential response to therapy. Ten models were selected for further evaluation, focusing on PDE3A and SLFN12 staining intensities in whole tissue sections. Finally, we selected two models for *in vivo* experiments: a leiomyosarcoma (UZLX-STS111) and a myxofibrosarcoma model (UZLX-STS274).

### *In vivo* synergy experiments

Female NMRI nu/nu mice were purchased from Janvier Laboratories (Le Genest-Saint-Isle, France). A total of 64 mice (32 mice per model) were used, aged 7–8 week-old and weighing 28–32 g at the time when PDX models were engrafted subcutaneously into both flanks. The two tumors in each animal were considered as independent events [24]. Prior to the start of the experiments, two mice from the UZLX-STS274 model were sacrificed due to lack of visible tumors, and one UZLX-STS111 mouse was found dead for unknown reasons. Tumors from the deceased UZLX-STS111 mouse were excluded from histological and molecular analysis.

Tumor volumes were determined in 31 mice engrafted with UZLX-STS111 and 30 with the UZLX-STS274 models using three digital caliper-measured dimensions (x, y, and z), calculated as tumor volume = (π × x × y × z)/6. Once half of the tumors reached a volume of 100 mm^3^, the mice were randomly assigned to one of 4 treatment groups. The treatment setup details, and dosing schedule are outlined in Table 1.

**Table 1.** Treatment and dosing schedule of the *in vivo* experiments.

| Group | Treatment | N of mice/tumors | N of mice/tumors | Morning treatment | Afternoon treatment |
| --- | --- | --- | --- | --- | --- |
| A | Vehicle control | 7 / 13 | 6 / 13 | 5 ml/kg vehicle of ANA | 5 ml/kg vehicle of A-1331852 |
| B | ANA | 8 / 13 | 7 / 14 | 10 mg/kg ANA | No treatment |
| C | A-1331852 | 8 / 15 | 8 / 15 | 25 mg/kg A-1331852 | No treatment |
| D | Combination | 8 / 14 | 8 / 15 | 10 mg/kg ANA | 25 mg/kg A-1331852 |

Abbreviations: ANA = anagrelide

The groups were treated as follows: (A) 5 ml/kg of both vehicles (ANA and A-1331852), (B) 10 mg/kg ANA in 10% ethanol, (C) 25 mg/kg A-1331852 in 60% (v/v) Phosal 50 PG (Cat. No. HY-Y1903, MedChemExpress), 27.5% (v/v) PEG400, 10% (v/v) ethanol, and 2.5% (v/v) DMSO, and (D) 10 mg/kg ANA and 25 mg/kg A-1331852. A-1331852 is an analog of A-1155463, with improved *in vivo* properties [27], and its vehicle formulation was prepared as previously described [28-34]. Mice were weighed daily to determine the administration volumes for the oral doses. Mice in groups B and C received a single daily treatment with ANA (10 mg/kg) and A-1331852 (25 mg/kg), respectively, as outlined in Table 1. The control group (A) received only the vehicles of ANA and A-1331852 at corresponding times. In the combination group (D), mice were first treated with ANA (10 mg/kg) in the morning, followed approximately 4 hours later by A-1331852 (25 mg/kg) in the afternoon. Body weights were measured regularly as an indicator of treatment tolerance, while animal well-being was visually monitored throughout the study. Tumor volumes were measured three times per week as a key indicator of treatment efficacy.

Mice were planned to be treated for four weeks. However, the treatment duration was shortened if significant tumor regression or lack of response was observed earlier, in order to adhere to ethical guidelines and minimize unnecessary animal distress. The UZLX-STS111 experiment concluded on day 19, and the UZLX-STS274 experiment on day 22. Mice were sacrificed with a lethal dose of pentobarbital (Dolethal, Vetoquinol, Lure, France), followed by cervical dislocation. Mice were weighed and blood samples were collected (described below). Tumors were collected and snap-frozen in liquid nitrogen. Tumors from mice that were euthanized before the end of the experiment were included in the histological assessment and IHC stainings. Organs were visually inspected during autopsy for signs of abnormalities, and livers were weighed and collected in 4% formaldehyde solution for later evaluation.

### Whole blood analysis

Blood samples were collected from the facial submandibular vein on day 0 after randomization into treatment groups, and via cardiac puncture following euthanasia at the end of the experiment. Samples were drawn into EDTA tubes and analyzed using the scil Vet abc Plus^+^ hematology analyzer (Scil Animal Care, Viernheim, Germany). Mice from which blood samples could not be collected at the end of the experiment due to premature death, were excluded from Paired-sample analyses.

### *In vivo* sample processing and evaluation

Formalin-fixed tumor and liver specimens were embedded in paraffin and sections of 3.5 μm were generated on regular microscope slides and stained with H&E for histological evaluation. Additionally, tumor FFPE sections were prepared on positively charged slides and IHC stained for Ki-67, PDE3A, and SLFN12. Human leukocyte antigen A (HLA-A) was additionally stained from 8 randomly selected tumors per model to prove human origin of the tumors. Staining conditions are shown in detail in Supplementary Table S1. PDE3A and SLFN12 staining intensities were scored on a scale of 0 to 3, while HLA-A was evaluated either positive or negative.

The proliferation rate was analyzed by calculating the average percentage of Ki-67-positive cells from 6 40x magnified images for each tumor. The ratio was calculated using the web-based ImmunoRatio 2.5 program [35] from the most representative tumor cell areas. Hthres and DABthres values were optimized for each slide before determining the percentage of positively stained cells [35]. For statistical analyzes, the average value for each tumor was used.

### Statistical analysis

Statistical analyses were performed using IBM SPSS Statistics 28.0 (Armonk, NY, USA). A significance level of 0.05 was used. Pearson correlation coefficient (*r*) was used to analyze correlation between combination DSRT results of ANA and BAY. Differences in *in vitro* relative cell viabilities across treatments were analyzed using one-way ANOVA and Tukey’s multiple comparison test. Tumor weights and relative tumor volumes, as well as initial blood component values between treatment groups were analyzed using the Independent-samples Kruskal–Wallis test and Dunn’s pairwise tests, adjusted with Bonferroni correction. Ki-67-positive cell ratios were analyzed with one-way ANOVA and Tukey’s multiple comparison test. Paired-samples Wilcoxon signed-rank test was used to investigate significant changes in blood component values during treatment.

## Results

### Discovering synergy between Bcl-2/Bcl-xL inhibitors and PDE3A modulators

We performed DSRT to analyze 527 drug combinations with two PDE3A modulators, ANA and BAY, in GIST882, SA4, and GOT3 sarcoma cell lines. The ΔDSS results of ANA and BAY (Supplementary Table S2) correlated across all three cell lines (GIST882: r = 0.698, *p* < 0.001; SA4: r = 0.716, *p* < 0.001; GOT3: r = 0.252, *p* < 0.001), indicating a similar mechanism of action of the drugs. PDE3A modulators had a smaller impact on compound toxicities in the GOT3 cell line compared to SA4 and GIST882. The top 30 synergistic and antagonistic compounds are presented in Figure 1a. PDE3A modulators increased DSS values in therapeutic drug classes targeting Bcl-2-like protein 2 (Bcl-xL) or both B-cell lymphoma 2 (Bcl-2) and Bcl-xL, as well as histone deacetylase (HDAC), mammalian target of rapamycin complex 1 (mTORC1), NEDD8-activating enzyme (NAE), p21 activated kinase (PAK), and aminopeptidase. Conversely, the PDE3A modulators acted as antagonists for many drugs, including several heat shock protein 90 (Hsp90) inhibitors, as well as those targeting bromodomain and extraterminal domain (BET) protein family, the MAPK/ERK pathway, and polo-like kinase 1 (PLK1). Interestingly, an antagonistic effect was seen with eribulin, a chemotherapeutic agent used in the treatment of breast cancer and liposarcoma [36].

**Figure 1.**
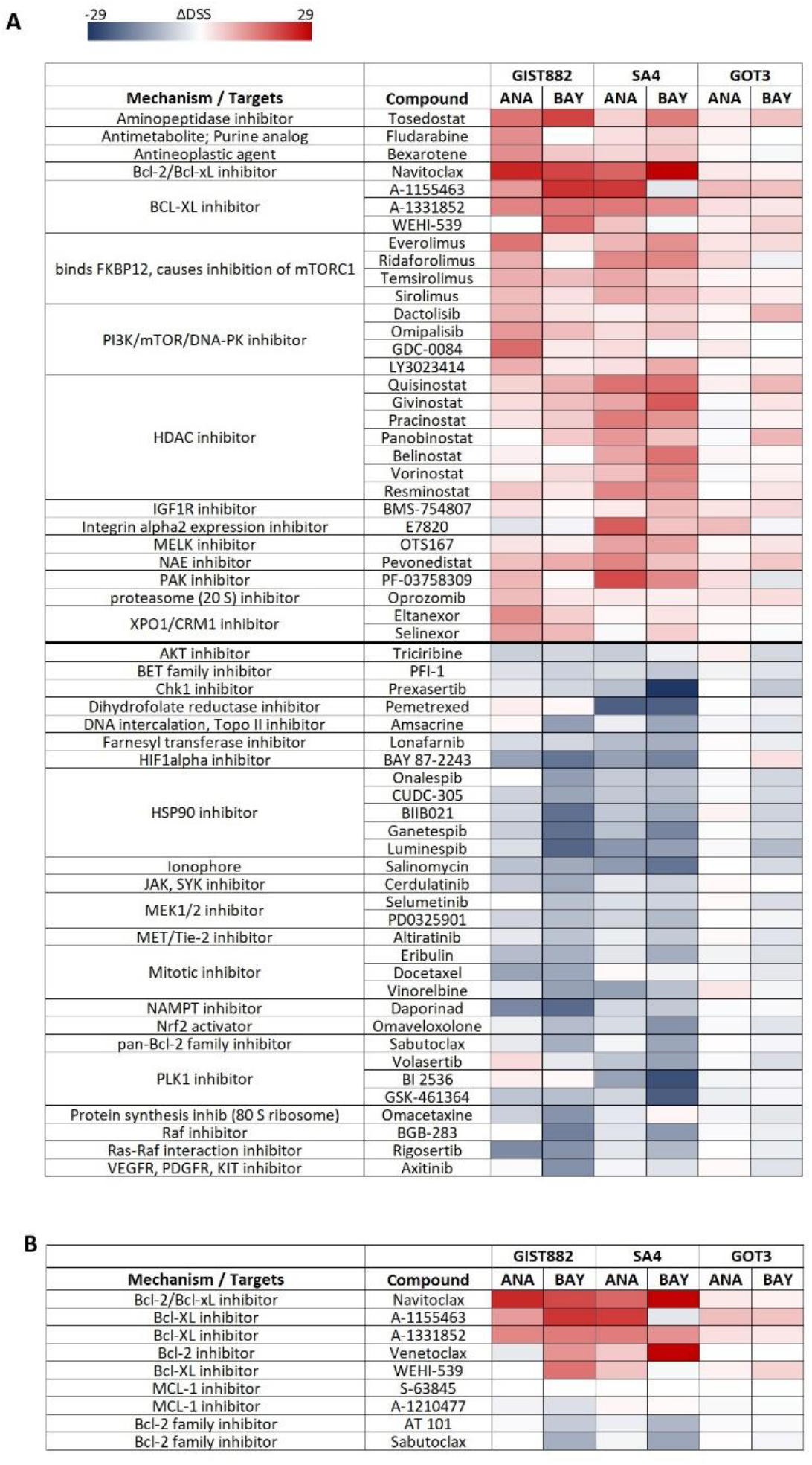
High-throughput drug screening identified compounds with altered efficacy in response to 100 nM ANA or 40 nM BAY in GIST882 and liposarcoma cell lines SA4 and GOT3. **(a)** The drug response heat map shows the ΔDSS results for the 30 most potentially synergistic combinations (in red) and the 30 most antagonistic combinations (in blue). **(b)** ANA and BAY showed synergy with Bcl-2/Bcl-xL and Bcl-xL targeting compounds in the cell lines.

Further investigation into inhibitors targeting the Bcl-2 family proteins (Figure 1b) revealed that PDE3A modulators specifically enhanced cytotoxic effects of inhibitors selectively targeting Bcl-xL and Bcl-2. In contrast, neither the Mcl-1 selective inhibitors, nor the pan-Bcl-2 family inhibitors demonstrated synergy with ANA or BAY; in fact, the latter indicated antagonism with BAY. Notably, the two pan-Bcl-2 family inhibitors, AT-101 and sabutoclax, both have a higher affinity for Mcl-1 compared to Bcl-2 and Bcl-xL [37, 38].

Based on the high-throughput drug combination results above, 10 compounds targeting Bcl-2/Bcl-xL, mTORC1, PAK, HDAC, and NAE were selected for examination in a 7×7 dose response matrix with ANA. Two independent experiments led to comparable results (Supplementary Table S3; Supplementary Figures S1–S3). Synergy strength was assessed using the ZIP model, and the average ZIP scores for drug combinations in each cell line are presented in Table 3. Notably, the two Bcl-xL-selective inhibitors, A-1155463 and A-1331852, and PAK inhibitor, PF-03758309, yielded high ZIP synergy scores across all three cell lines. Considering the synergy concentration distributions (Supplementary Figures S1–S3), 100 nM of ANA, A-1155463, and navitoclax were chosen for the subsequent cell viability experiments.

**Table 3.**
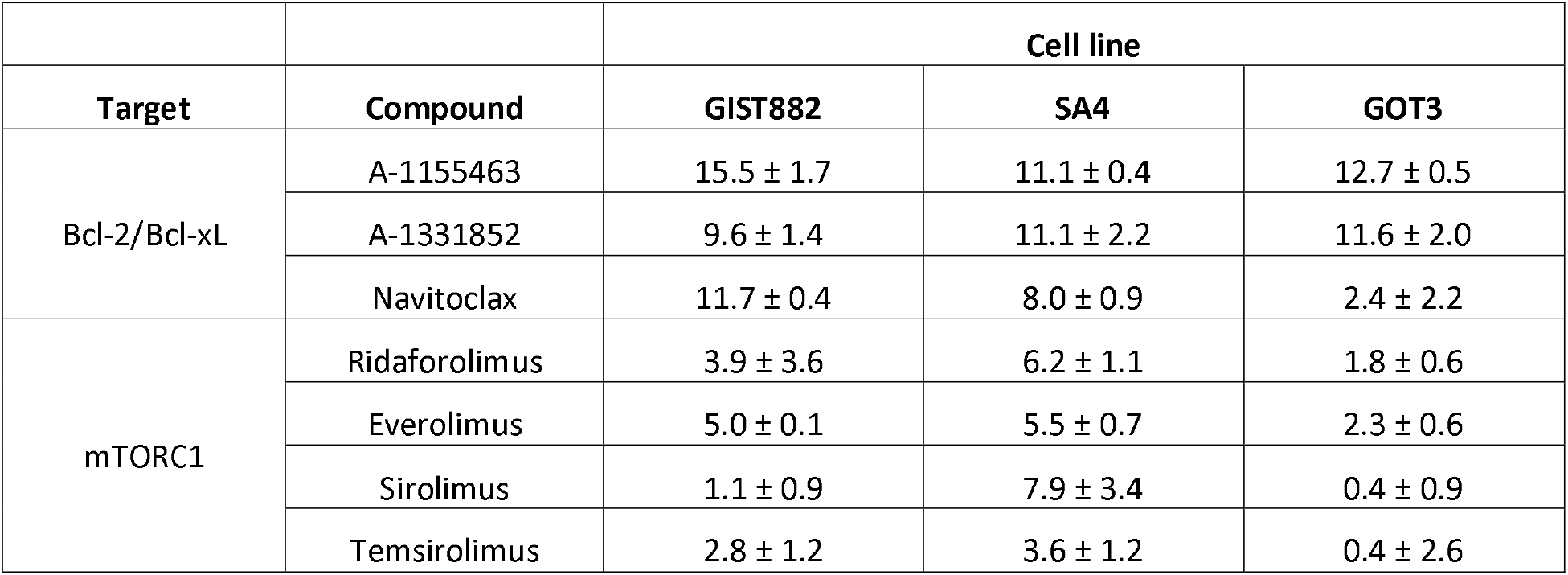
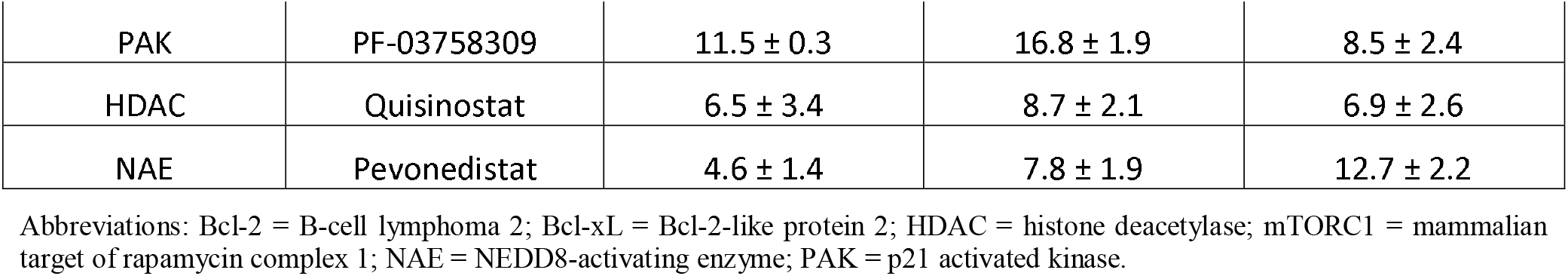
Synergy between selected compounds and ANA was assessed with ZIP model in three sarcoma cell lines. Data are presented as mean ± standard deviation (SD) of two separate experiments.

| Target | Compound | Cell line |  |  |
| --- | --- | --- | --- | --- |
|  |  | GIST882 | SA4 | GOT3 |
| Bcl-2/Bcl-xL | A-1155463 | 15.5 ± 1.7 | 11.1 ± 0.4 | 12.7 ± 0.5 |
| | A-1331852 | 9.6 ± 1.4 | 11.1 ± 2.2 | 11.6 ± 2.0 |
| | Navitoclax | 11.7 ± 0.4 | 8.0 ± 0.9 | 2.4 ± 2.2 |
| mTORC1 | Ridaforolimus | 3.9 ± 3.6 | 6.2 ± 1.1 | 1.8 ± 0.6 |
| | Everolimus | 5.0 ± 0.1 | 5.5 ± 0.7 | 2.3 ± 0.6 |
| | Sirolimus | 1.1 ± 0.9 | 7.9 ± 3.4 | 0.4 ± 0.9 |
| | Temsirolimus | 2.8 ± 1.2 | 3.6 ± 1.2 | 0.4 ± 2.6 |
| PAK | PF-03758309 | 11.5 ± 0.3 | 16.8 ± 1.9 | 8.5 ± 2.4 |
| HDAC | Quisinostat | 6.5 ± 3.4 | 8.7 ± 2.1 | 6.9 ± 2.6 |
| NAE | Pevonedistat | 4.6 ± 1.4 | 7.8 ± 1.9 | 12.7 ± 2.2 |

Abbreviations: Bcl-2 = B-cell lymphoma 2; Bcl-xL = Bcl-2-like protein 2; HDAC = histone deacetylase; mTORC1 = mammalian target of rapamycin complex 1; NAE = NEDD8-activating enzyme; PAK = p21 activated kinase.

### PDE3A modulators sensitize cancer cells to Bcl-2/-xL inhibitors

Following the observed synergy between Bcl-2/-xL inhibitors and PDE3A modulators, we expanded our research to include DNMDP (6-(4-(diethylamino)-3-nitrophenyl)-5-methyl-4,5-dihydropyridazin-3(2H)-one), a well-established PDE3A modulator, for more comprehensive evaluation. A 10 nM concentration was chosen for DNMDP as it approximately corresponded to the 100 nM ANA concentration (Supplementary Figure S4). In addition to the cell lines above, we included three more—the neuroglioma cell line H4, and melanoma cell lines CHL-1 and A2058—to analyze the synergy of Bcl-2/-xL inhibitors, navitoclax and A-1155463, with PDE3A modulators, ANA and DNMDP. All cell lines co-expressed PDE3A and SLFN12 (Figure 2a). On their own, ANA and DNMDP significantly reduced cell viability across all cell lines (Figure 2b; Supplementary Figure S5). Both PDE3A modulators sensitized GIST882, A2058, H4, and CHL-1 cell lines to the Bcl-2/Bcl-xL inhibitors, enhancing treatment efficacy. The Bcl-2/Bcl-xL inhibitors did exhibit no cytotoxic effects alone in any of the cell lines, with the exception of A-1155463, which demonstrated activity in the SA4 and GOT3 cell lines.

**Figure 2.**
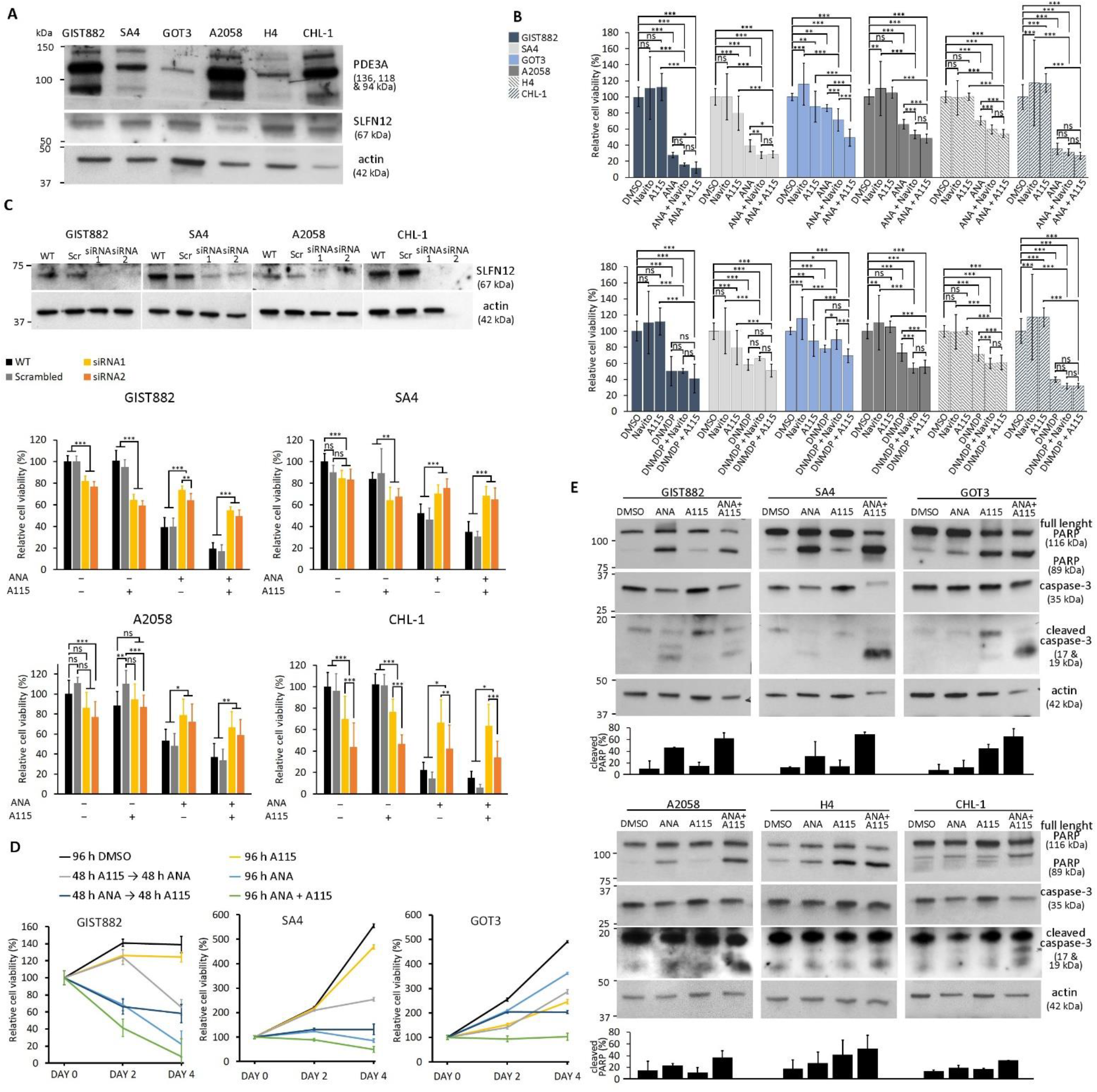
PDE3A modulators increase the sensitivity of tumor cell lines to Bcl-2/-xL inhibitors. **(a)** PDE3A and SLFN12 expression was detected in six cell lines. **(b)** The combination of ANA or DNMDP with two Bcl-2/-xL inhibitors, navitoclax and A-1155463, was investigated in the cell lines. The combination treatments were most effective in reducing cell viability after 72 hours. Data are presented as mean ± standard deviation from two independent experiments. Statistical significance was determined by one-way ANOVA with Tukey’s multiple comparisons test. **(c)** A siRNA-mediated SLFN12 knockdown (above) attenuated the efficacy of ANA and the synergistic combination in all tested cell lines. Data are shown as mean ± standard deviation from two independent experiments, analyzed using one-way ANOVA with Tukey’s test. **(d)** Simultaneous treatment with ANA and A-1155463 resulted in the greatest reduction in cell viability compared to sequential treatments. Data are presented as mean ± standard deviation from two independent experiments. **(e)** The combination of ANA and A-1155463 enhanced PARP and caspase-3 cleavage. Densitometry data (below) are from two separate western blots presented as mean ± standard deviation. Abbreviations: A115 = A-1155463; ANA = anagrelide; Navito = navitoclax. \**p* < 0.05, \*\**p* < 0.01, \*\*\**p* < 0.001, ns = not significant

### The synergy mechanism depends on the PDE3A-SLFN12 complex formation

To investigate whether synergy depends on PDE3A modulator-induced PDE3A-SLFN12 complex formation, we knocked down SLFN12 protein expression in GIST882, SA4, A-2058, and CHL-1 cell lines and treated cells with ANA or A-1155463, or both, and monitored subsequent cell viability (Figure 2c; Supplementary Figure S6). Indeed, the absence of SLFN12 diminished efficacy of ANA and combination treatment. The combination of ANA and A-1155463 was also tested in cell lines unresponsive to PDE3A modulators: GIST-T1, GIST48, and MLS402-91. GIST-T1 lacked SLFN12 expression and liposarcoma cell line MLS402-91 lacked PDE3A expression, likely explaining their insensitivity to the modulators (Supplementary Figure S7a). Insensitivity of PDE3A and SLFN12 positive GIST48 cell line has been reported earlier [14]. No synergy between ANA and A-1155463 was observed in any of the cell lines (Supplementary Figure S7b). In GIST48, the A-1155463 alone reduced cell viability, and the addition of ANA failed to enhance this reduction.

The combination of A-1155463 and the PDE3A inhibitor cilostazol did not demonstrate synergistic effects (Supplementary Figure S7c), suggesting that the previously observed sensitizing effect is independent of PDE3A enzyme inhibition. Notable exceptions were the GIST882 and GOT3 cell lines, where the combination reduced cell viability compared to cilostazol alone. However, in GOT3, A-1155463 on its own was cytotoxic, and in GIST882 cells, the combination with cilostazol was still less cytotoxic than ANA. Taken together, these results suggest that the synergistic effect is mediated by the formation of the PDE3A-SLFN12 complex.

Next, we investigated whether simultaneous drug exposure is necessary to achieve synergy, or if alternating administration could yield similar effects. Comparing sequential and simultaneous exposure of ANA and A-1155463 in GIST882, SA4, and GOT3 cell lines showed that simultaneous treatment was required to induce synergy (Figure 2d).

### Insights into ANA- and synergy-induced apoptosis mechanisms

Activation of apoptosis was assessed by evaluating PARP and caspase-3 cleavage in both single and combination treatments across PDE3A modulator-sensitive cell lines (Figure 2e). PARP cleavage was most prominent following the combination treatment in all cell lines, while caspase-3 cleavage was observed in SA4, GOT3, A2058, and CHL-1 cells. A-1155463 on its own induced PARP cleavage in GOT3 and H4 cells. Previous studies have reported that PDE3A modulators reduce expression levels of anti-apoptotic Bcl-2 family proteins, including Bcl-2, Bcl-xL, and Mcl-1 [3, 4, 17]. However, we observed no consistent changes in the expression of these proteins across all cell lines (Supplementary Figure S8).

To investigate how ANA affects apoptosis-related signaling pathways, which may explain its sensitizing effect to Bcl-xL and Bcl-xL/-2 inhibitors, we performed TaqMan Array Human Apoptosis analysis on ANA-treated cells. After successful RT-qPCR on GIST882, SA4, GOT3, A2058, and CHL-1 cell lines, we evaluated gene expression fold changes induced by a 24-hour ANA treatment (Supplementary Figure S9). Although no specific group of apoptotic markers was consistently upregulated or downregulated across all cell lines, several notable individual genes emerged.

In all 5 cell lines, ANA increased the expression of *TNFRSF10B* and *BBC3*, whose corresponding proteins are known to be upregulated by PDE3A modulators [16, 17]. *TNFSF10* was also one of the highly upregulated genes across all cell lines, with its translated protein, TRAIL, described as synergizing with ANA [17]. Other genes consistently upregulated in all 5 cell lines included *BCL2L13, BCL2L2, BNIP3L, BOK, NOD1, CASP1, CASP10, CASP4, CASP8, CASP9, CFLAR, CRADD, IKBKB, LRDD*, and *PMAIP1*. In contrast, *BIRC5, CHUK, NFKBIA, TNFRSF21*, and *BID* were downregulated in most of the cell lines.

No significant downregulation of Bcl-2 family genes was observed across the cell lines, and the results for *BCL2* and *MCL1* varied between cell lines. *BCL2* was downregulated only in GIST882 and CHL-1, and *MCL1* only in A2058. Interestingly, *BCL2L1* (Bcl-xL) was upregulated in all cell lines except GIST882.

### ANA and A-1331852 synergized *in vivo* in a PDE3A and SLFN12 double positive sarcoma PDX model

PDX models from 9 different sarcoma subtypes were grouped by PDE3A and SLFN12 expression, as detected by IHC on TMAs. As sensitivity to PDE3A modulators correlates with the co-expression of PDE3A and SLFN12 [2, 10, 12-17], models with staining intensities of ≥1 for PDE3A and ≥0.25 for SLFN12 were considered potentially sensitive to treatment. The lower intensity threshold for SLFN12 was determined based on our previous study [39], which showed that only faint SLFN12 staining by IHC was sufficient to produce high efficacy of ANA in a GIST PDX model UZLX-GIST2B. Double positivity for both markers was observed in 32% (23/73) of sarcoma PDXs (Figure 3a; Supplementary Table S4). After confirming the results using IHC on ten selected whole tissue sections from xenograft tissue, two models—leiomyosarcoma UZLX-STS111 and myxofibrosarcoma UZLX-STS274—were selected for the following *in vivo* synergy experiments (Figure 3b).

**Figure 3.**
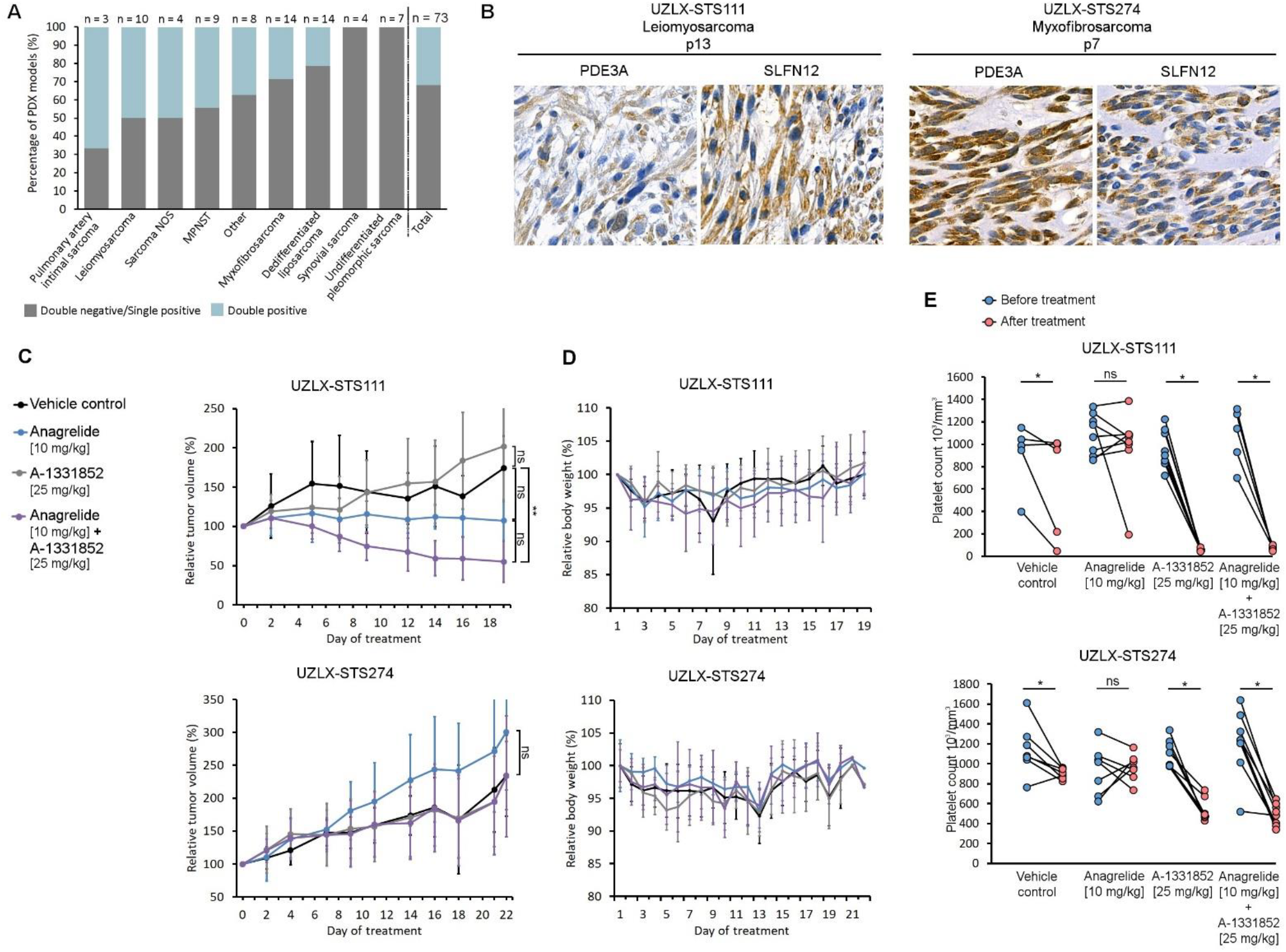
Synergistic effects of ANA and A-1331852 in sarcoma PDX mouse models. **(a)** 23 of 73 sarcoma PDX model tissues were double positive for PDE3A and SLFN12. **(b)** The two selected PDX models, UZLX-STS111 and UZLX-STS274, showed PDE3A and SLFN12 expression by IHC. **(c)** The leiomyosarcoma UZLX-STS111 model showed a significant reduction in tumor volume in response to the drug combination therapy (Independent-Samples Kruskal-Wallis Test), whereas the UZLX-STS274 model did not. **(d)** No significant alterations in body weight were observed in the mice across any of the treatment groups. **(e)** Mice receiving A-1331852 had a significant reduction of platelet count during the treatment (Paired-samples Wilcoxon signed-rank test). Abbreviations: MPNST = Malignant peripheral nerve sheath tumor; NOS = Not otherwise specified; p = passage. \**p* < 0.05; \*\**p* < 0.001; ns = not significant.

Mice were treated by oral gavage with 10 mg/kg ANA, 25 mg/kg A-1331852, or a combination of both. The combination treatment demonstrated anticancer activity in the leiomyosarcoma model, UZLX-STS111, significantly reducing tumor volumes (Figure 3c; Independent Samples Kruskal-Wallis test, *p* < 0.001), while ANA indicated stabilization of tumor growth. Tumors treated with the combination therapy also weighed the least at time of collection at the end of the experiment (Supplementary Figure S10). A-1331852 alone showed no antitumor efficacy. None of the treatments significantly affected tumor volumes or weights in the myxofibrosarcoma model, UZLX-STS274.

The treatments were generally well tolerated, with no significant weight loss of mice observed (Figure 3d). However, one vehicle control mouse from the UZLX-STS111 model was sacrificed due to over 20% weight loss, and two mice from the combination treatment group died immediately after oral gavage. Additionally, one vehicle control mouse from UZLX-STS111 died overnight. All of these mice were from groups receiving twice-daily gavage, which appeared to cause more agitation compared to the single-agent groups. Early in the experiments, all mice receiving A-1331852 either alone or with ANA, exhibited mild skin erythema, which resolved within a week (Supplementary Figure S11a). Dehydrated skin was also frequently observed in mice from these treatment groups. Importantly, no hepatotoxicity was observed in any mouse (Supplementary Figure S11b).

Although tumor regression was seen in the combination-treated UZLX-STS111 tumors, histopathological evaluation revealed no differences in histology or in the ratio of Ki-67-expressing tumor cells between treatment groups in either PDX model (Figure 4; Supplementary Figure S12). Notably, SLFN12 expression in UZLX-STS274 tumors was significantly lower than observed during model selection, with some tumors showing no expression or only faint staining. PDE3A and SLFN12 expression levels remained unchanged across all treatment groups suggesting that none of the treatments affected their protein expression. The tumors were confirmed to be of human origin by HLA-A positivity.

**Figure 4.**
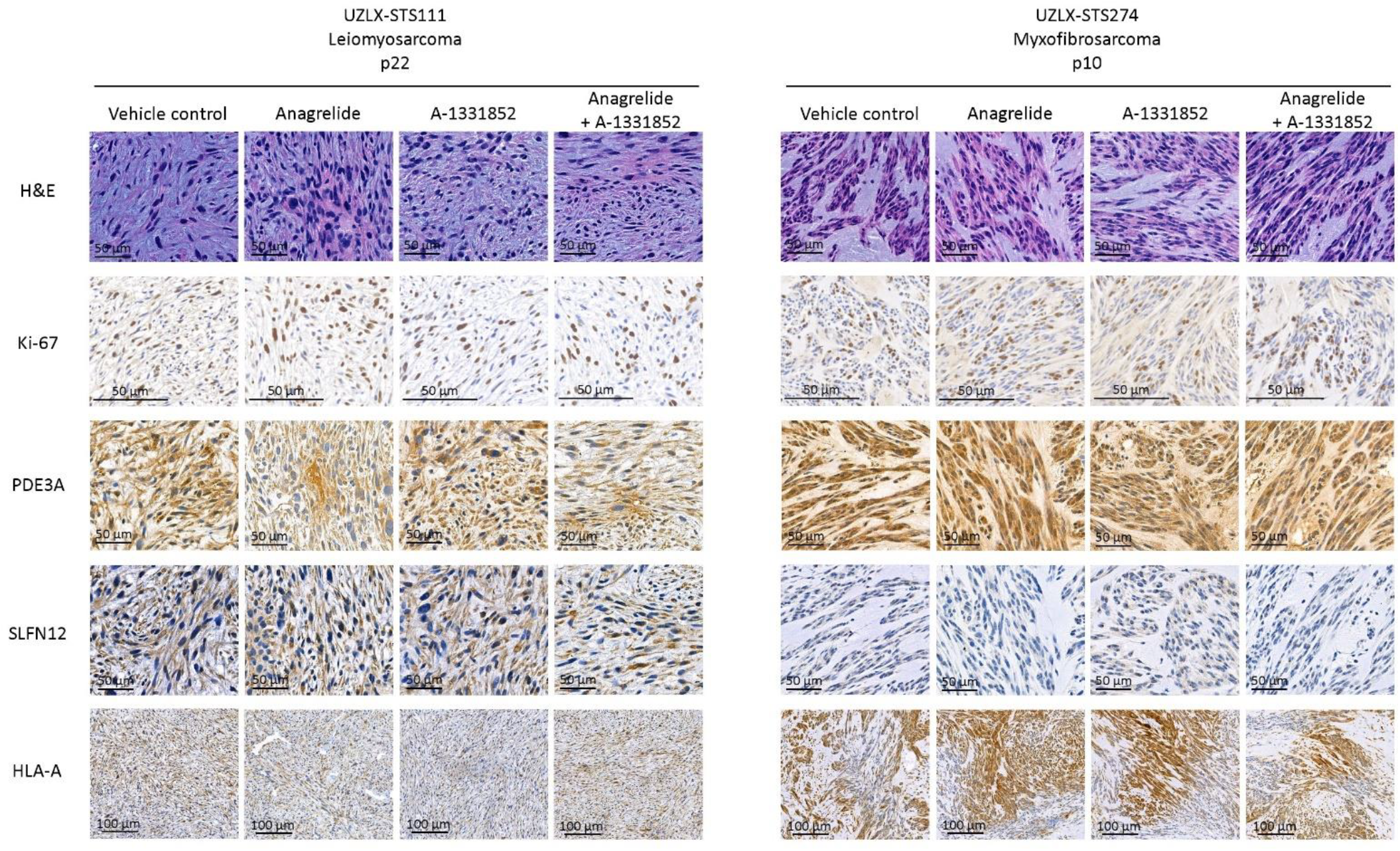
*In vivo* PDX tumors were histologically and molecularly characterized.

### A-1331852 affected hematological parameters more than ANA

Blood samples collected before and after the experiments were analyzed using a hematology analyzer to evaluate any changes in blood parameters. The results were consistent across both PDX models (Supplementary Figure S13). Additionally, blood parameters were similar across treatment groups in both PDX models before starting the treatments (Independent-samples Kruskal–Wallis test, *p* > 0.05). In both experiments, A-1331852, either alone or in combination with ANA, significantly decreased platelet counts (Figure 3e, Paired-samples Wilcoxon signed-rank test, *p* < 0.05), while ANA alone did not. Furthermore, red blood cell counts, hemoglobin, and hematocrit decreased, while mean corpuscular volume, mean corpuscular hemoglobin, red blood cell distribution width, and platelet volumes increased (Paired-samples Wilcoxon signed-rank test, *p* < 0.05) in mice receiving A-1331852, either alone or in combination. Splenomegaly was observed in 8 of 26 UZLX-STS111 mice and 2 of 30 UZLX-STS274 mice, all of which received A-1331852, either as a single agent or in combination with ANA.

## Discussion

High-throughput drug screening studies have been widely and successfully employed to identify synergistic combinations in oncology [40-42]. Utilizing this approach, we discovered synergistic compounds that enhance treatment efficacy in PDE3A expressing tumors. Such supportive combinatorial compounds can expand therapeutic options and help optimize dosing strategies. PDE3A modulators demonstrated synergy in cell lines with inhibitors targeting Bcl-2/Bcl-xL, mTORC1, PAK, HDAC, and NAE. Focusing on inhibitors that target anti-apoptotic proteins of the Bcl-2 family, we found that PDE3A modulators sensitize cancer cells to Bcl-2/Bcl-xL inhibitors but do not influence responses to Mcl-1-selective or pan-Bcl-2 family inhibitors. As one of the cancer hallmarks is evasion of apoptosis, overexpression of the antiapoptotic Bcl-2 family proteins can contribute to this feature in some cancer types [43, 44]. Indeed, selective inhibitors targeting the Bcl-2 family have been assessed in clinical trials [45-47]. While Bcl-xL-selective inhibitors are yet to be investigated, existing evidence suggests that they may impact on the human hematopoietic system [48].

We investigated the mechanism of action of PDE3A modulators and their ability to sensitize cells to Bcl-2/Bcl-xL inhibitors. Synergy was not observed after SLFN12 was knocked down, in PDE3A modulator-unresponsive cell lines, or with a PDE3A enzyme inhibitor. Given that PDE3A and SLFN12 are not known to interact with each other otherwise [3-5], these findings suggest that the PDE3A-SLFN12 complex is crucial for synergy. Prior studies have reported that the PDE3A-SLFN12 complex disrupts mitochondrial membrane potential [3], and that mitochondrial membrane damage improves target access of A-1331852 [49]. The synergy between PDE3A modulators and Bcl-xL inhibitors could operate through a similar mechanism, even though further investigation is needed to confirm this. The observed increase in apoptosis, recognized by PARP and caspase-3 cleavage, along with the greater efficacy of simultaneous versus sequential exposure, would certainly align with this hypothesis.

When investigating the apoptotic signaling pathways induced by ANA, some inconsistencies with prior findings emerged [3, 17]. *BCL2, BCL2L1*, and *MCL1* were not clearly downregulated across all cell lines, as confirmed by Western blot analysis. Notably, *BCL2L1*, which encodes both the pro-survival protein Bcl-xL and the pro-apoptotic protein Bcl-xS through alternative splicing in exon 1 [50], was elevated in all cell lines except GIST882. Bcl-xS interacts with Bcl-xL and Bcl-2 [51], and activates the pro-apoptotic proteins, Bak and Bax, leading to cytochrome c release [52]. Whether ANA promotes apoptosis by favoring alternative splicing towards the pro-apoptotic Bcl-xS, or if cells upregulate Bcl-xL expression as a survival mechanism in response to ANA exposure, remains to be investigated.

Endoplasmic reticulum stress activates the eIF2a/ATF4 signaling pathway [53]. The same pathway has also been shown to be activated during ANA-induced inhibition of megakaryocyte differentiation, leading to reduced platelet counts [54-56]. Additionally, a PDE3A modulator, OPB-171775, was reported to increase ATF4 expression in primitive neuroectodermal tumor cells and a GIST PDX model [16], while DNMDP exposure induced phosphorylation of GCN2, which subsequently phosphorylates eIF2α [57]. The pathway is known to upregulate *BBC3, TNFRSF10B, PMAIP1*, and *CDKN1A* (p21) [55], of which the first three were elevated in our TaqMan array, while p21 has previously been described as upregulated following PDE3A modulator treatment [16, 17, 19]. The eIF2α/ATF4 pathway is associated with HDAC [55], mTOR [58], the MAPK/ERK pathway [59], Hsp90 [56], and HIF-1α [60]. Interestingly, several inhibitors targeting these pathways emerged in our combination DSRT as synergistic or antagonistic, although the use of only three sarcoma cell lines may limit the specificity of the findings. The possibility that PDE3A modulators exert cytotoxic effects through activation of the eIF2α/ATF4 pathway is intriguing and warrants further investigation.

Finally, we evaluated the combination treatment *in vivo* in two sarcoma PDX models, using positivity for both PDE3A and SLFN12 expression as a basis for classification. PDE3A positivity was frequently observed in the leiomyosarcoma models, consistent with previous studies [9, 14]. Half of these models were also double positive for expression, highlighting the potential of PDE3A modulators as a targeted therapy. Indeed, we observed drug synergy in the UZLX-STS111 leiomyosarcoma model, where ANA stabilized tumor growth, while the combination significantly reduced tumor size. The other selected PDX model, UZLX-STS274, showed decreased SLFN12 expression intensity during tumor tissue passaging in mice, which may explain its lack of response to therapy. Since protein expression can change during passaging [61-63], evaluating drug target levels in PDX tissue from closely relevant passages is essential. Investigating whether SLFN12 expression levels are influenced by changes in interferon levels and the tumor microenvironment in immunocompromised mice could provide valuable insights, as interferons have been shown to elevate SLFN12 expression [17]. However, in our previous study, we demonstrated that even low SLFN12 expression can yield strong responses to ANA in a GIST PDX model [39]. This may have been due to ANA’s stabilizing effect on SLFN12, potentially leading to increased SLFN12 expression [3, 5, 16] and heightened sensitivity to treatment. Furthermore, mutations in PDE3A or SLFN12 that affect complex formation should be considered, as they have been linked to resistance [4, 5, 15].

Tolerability of the *in vivo* treatments was also assessed. The treatments did not significantly affect mouse body weight, and A-1331852 combined with ANA did not cause acute liver toxicity, unlike when combined with the MCL-1 inhibitor S63845 [30] or the CDK inhibitor Dinaciclib [32]. However, A-1331852 had a greater impact on blood parameters than ANA. In addition to the decreased platelet counts previously associated with A-1331852 [28, 29, 34], mice treated with A-1331852 exhibited increased mean corpuscular volume and mean corpuscular hemoglobin concentration values, along with reduced hematocrit and platelet counts, and enlarged spleens. Similar changes have been reported in *BCL2L1*-deficient mice, where enlarged spleens were attributed to accelerated erythropoiesis in response to anemia [64]. ANA is clinically used to treat a condition, in which a patient has elevated platelet counts. It inhibits megakaryocyte differentiation and is suggested to do so independently from PDE3A enzyme inhibition [1, 65, 66]. Since mice lack a direct ortholog of human SLFN12 [67] and we observed no decrease in platelet count in the animals, it is reasonable to hypothesize that ANA reduces platelet levels through the PDE3A-SLFN12 complex. Similar results have been reported in dogs [68]. Until recently, ANA was the only PDE3A modulator used in humans. Similar findings—reduced platelet counts in humans but not in rodents—were reported with BAY [69], suggesting a consistent mechanism across PDE3A modulators.

In addition to the oncological opportunities, the synergistic approach may also benefit patients with refractory essential thrombocythemia. Although ANA achieves a good response rate of approximately 80% in essential thrombocythemia patients [70, 71], synergistic drug combinations could offer broader disease management. This approach could achieve the same therapeutic effect with lower doses or reduce platelet counts more rapidly and to lower levels in patients with extremely high platelet counts.

In this study, we identified synergy between PDE3A modulators and multiple targeted agents investigated for oncological purposes. Since many cancer subtypes still lack effective clinical strategies, novel treatment alternatives that specifically utilize precision medicine are needed. PDE3A is an emerging therapeutic target and PDE3A modulators show potential for treating soft-tissue sarcomas, tyrosine kinase inhibitor-resistant GIST, and ovarian cancer. Future studies should evaluate PDE3A and SLFN12 expression as key indicators of sensitivity to PDE3A modulators and explore other potential predictive biomarkers for treatment sensitivity. Additionally, the clinical potential of synergistic treatments warrants further investigation.

## Abbreviations

ANA: Anagrelide
BAY: BAY 2666605
Bcl-2: B-cell lymphoma 2
Bcl-xL: Bcl-2-like protein 2
BET: bromodomain and extraterminal domain
DSRT: Drug sensitivity and resistance tests
HDAC: Histone deacetylase
HLA-A: Human leukocyte antigen A
Hsp90: Heat shock protein 90
IHC: Immunohistochemistry
mTORC1: Mammalian target of rapamycin complex 1
NAE: NEDD8-activating enzyme
PAK: p21 activated kinase
PDE3A: Phosphodiesterase 3A
PDX: Patient-derived xenograft
PLK: polo-like kinase 1
SLFN12: Schlafen 12

## Data availability

Data are available from the corresponding author upon reasonable requests.

## Acknowledgements

The DSRT studies were carried out at the FIMM High Throughput Biomedicine Unit, which is hosted by the University of Helsinki and supported by HiLIFE and Biocenter Finland. The *in vivo* PDX experiments were conducted at KU Leuven, where the sarcoma PDX TMAs were also constructed and obtained. The study was funded by grants from the Jane and Aatos Erkko Foundation, the Sarcoma Foundation of America, the University of Helsinki, the Liv och Hälsa Foundation, the Relander Foundation, the Finnish Society of Sciences and Letters, the Avohoidon Tutkimussäätiö Foundation, the Finska Läkaresällskapet foundation, the Emil Aaltonen Foundation, the Academy of Finland, and the Sigrid Juselius foundation. We thank Biocenter Finland-supported FIMM High-Throughput Biomedicine Unit for providing pre-plated drug plates and FIMM Sequencing Unit for providing the sequencing services. We thank Philipp Ianevski for drug synergy analysis.

## Author contributions

**K. Toivanen:** Investigation, project administration, validation, visualization, writing–original draft, formal analysis, conceptualization. **A. Murumägi:** Resources, writing–review & editing. **A. Wozniak:** Resources, investigation, supervision, writing–review & editing. **K. Wyns:** Investigation, writing–review & editing. **C.-C. Wang:** Investigation, writing–review & editing. **L. De Sutter:** Investigation, writing–review & editing. **M. Arjama:** Investigation, resources, writing– review & editing. **N. Merikoski:** Investigation, writing–review & editing. **S. Salmikangas:** Investigation, writing–review & editing. **O. Youssef:** Investigation, writing–review & editing. **J. Isola:** Investigation, writing–review & editing. **O. Kallioniemi:** Resources, writing–review & editing. **P. Schöffski:** Resources, investigation, writing–review & editing. **T. Böhling:** Resources, supervision, writing–review & editing, funding acquisition, conceptualization. **H. Sihto:** Project administration, supervision, writing–original draft, funding acquisition, conceptualization.

