## Supplementary Figures for "Phosphodiesterase 3A modulators sensitize tumor cells to Bcl-xL and Bcl-2/Bcl-xL inhibitors"

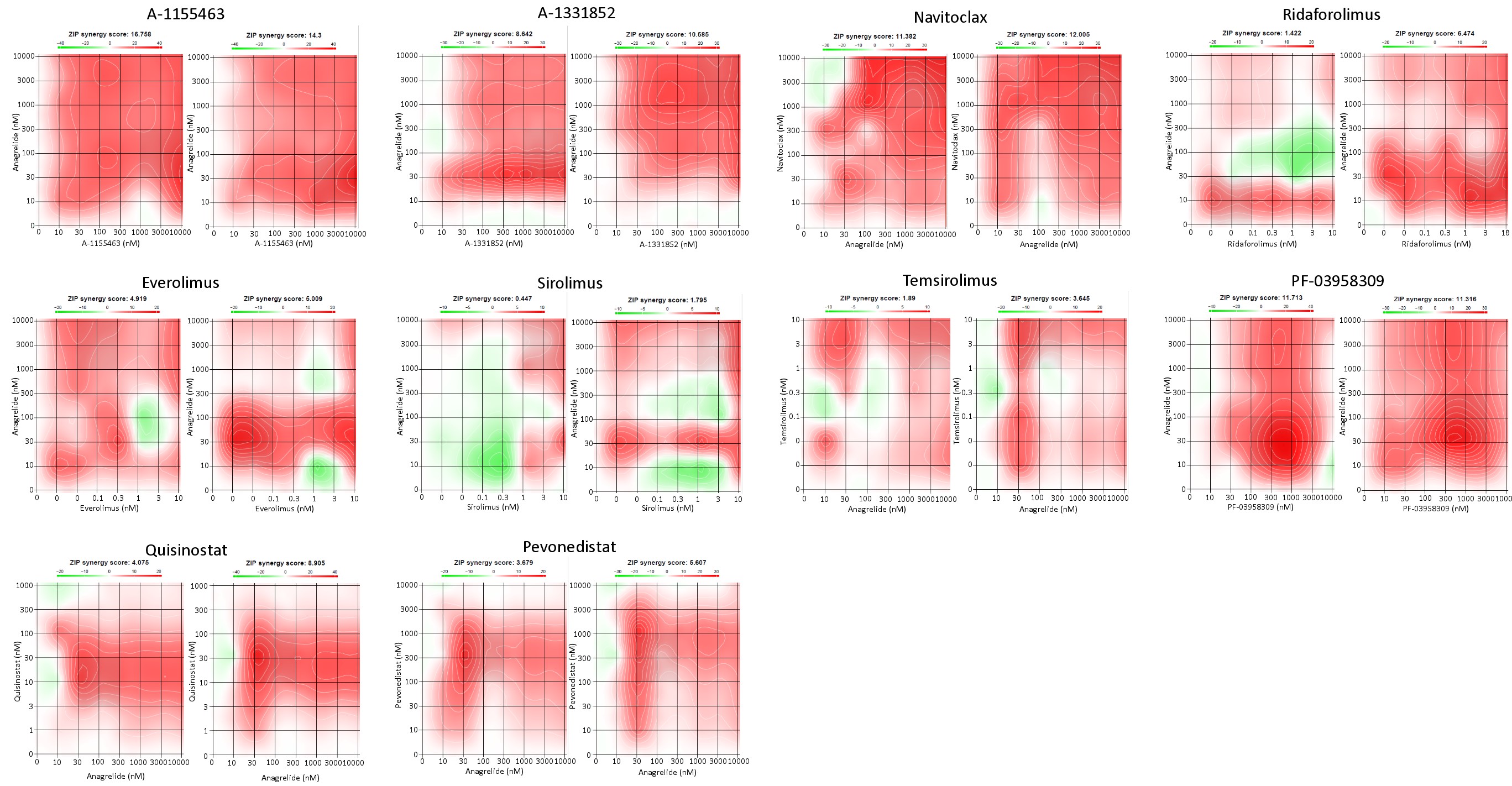

**Figure S1**. Visualization of dose-response matrix between anagrelide and selected 10 compounds in GIST882 cell line.

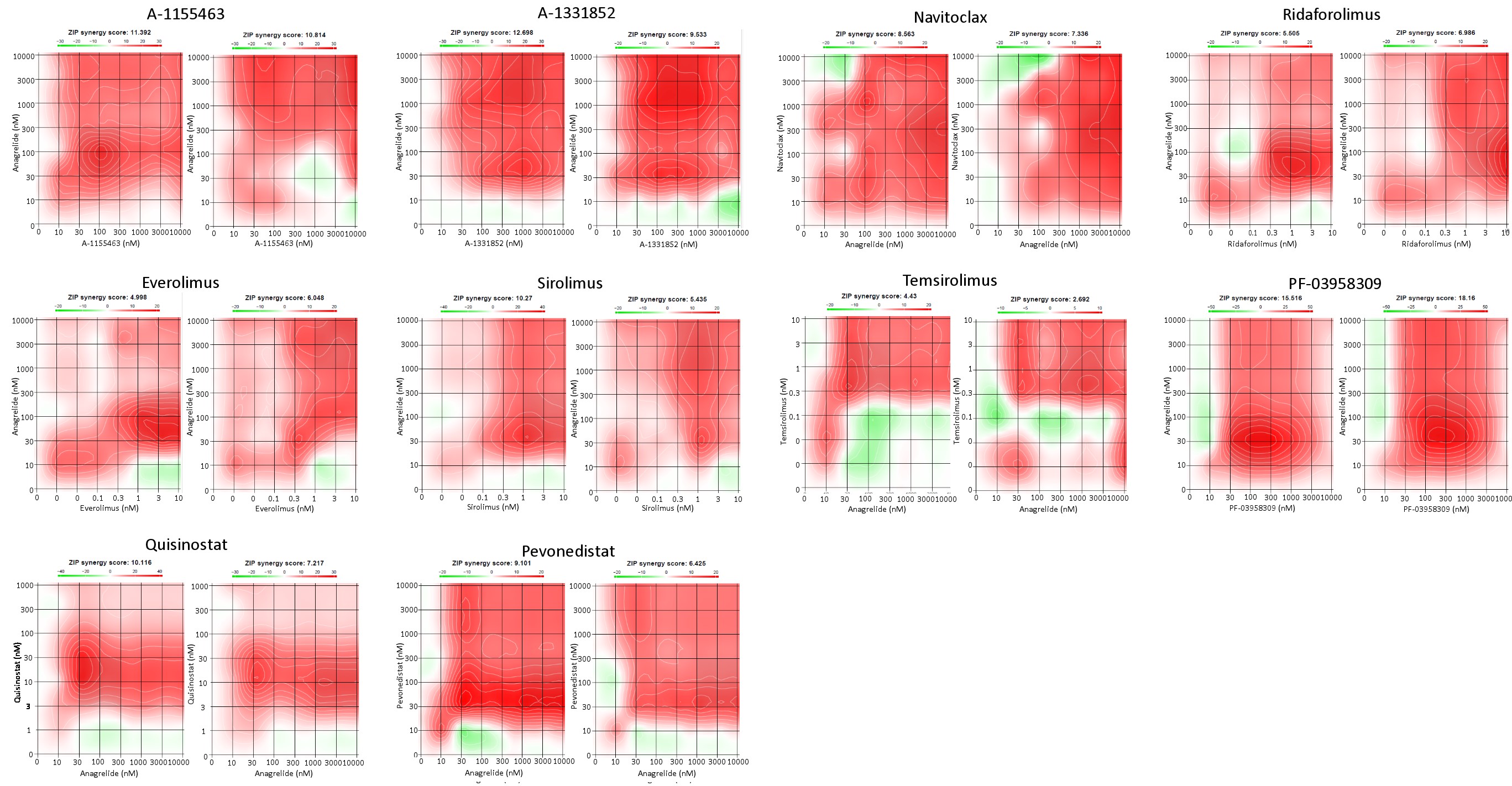

**Figure S2**. Visualization of dose-response matrix between anagrelide and selected 10 compounds in SA4 cell line.

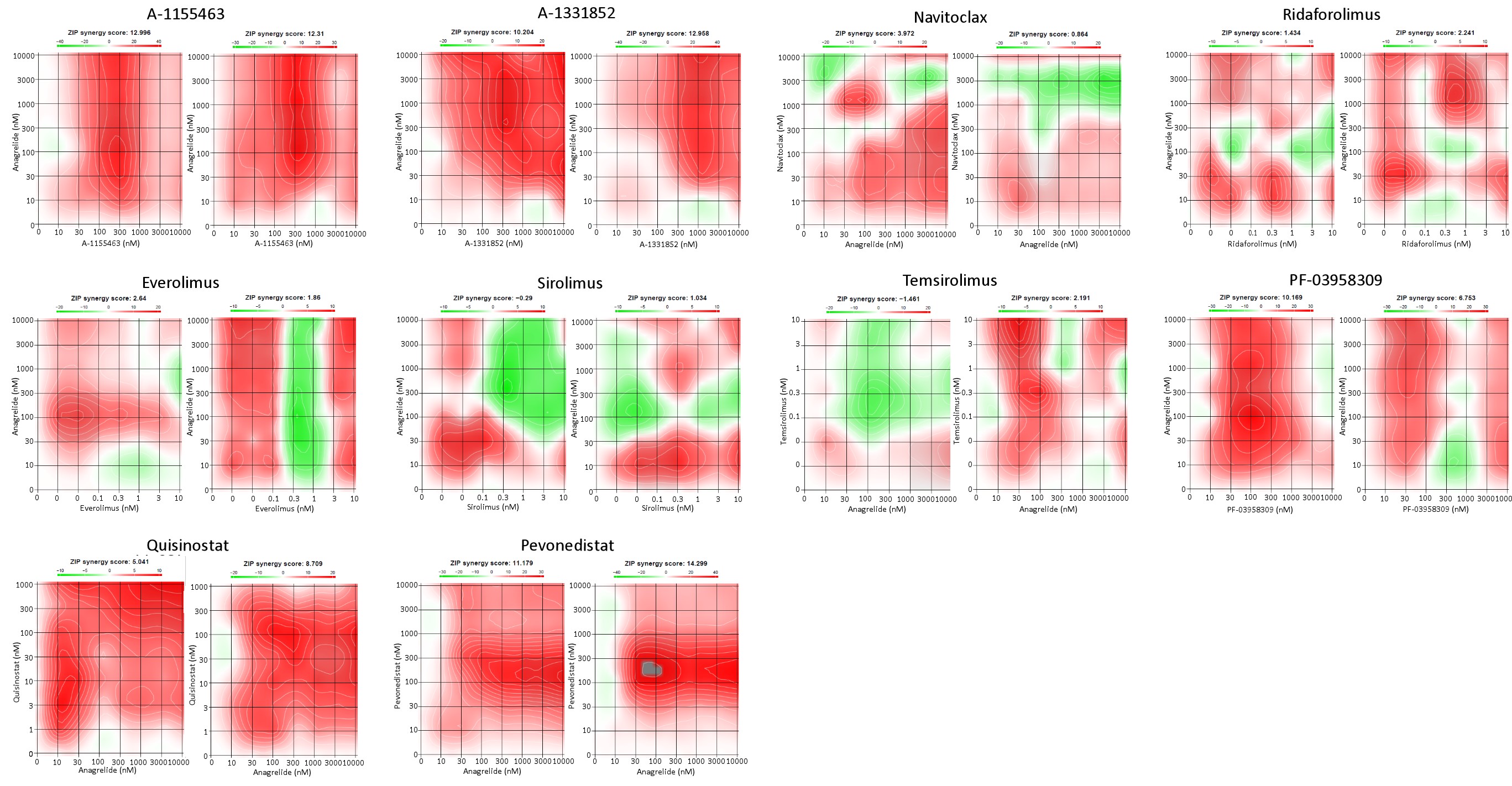

**Figure S3**. Visualization of dose-response matrix between anagrelide and selected 10 compounds in GOT3 cell line.

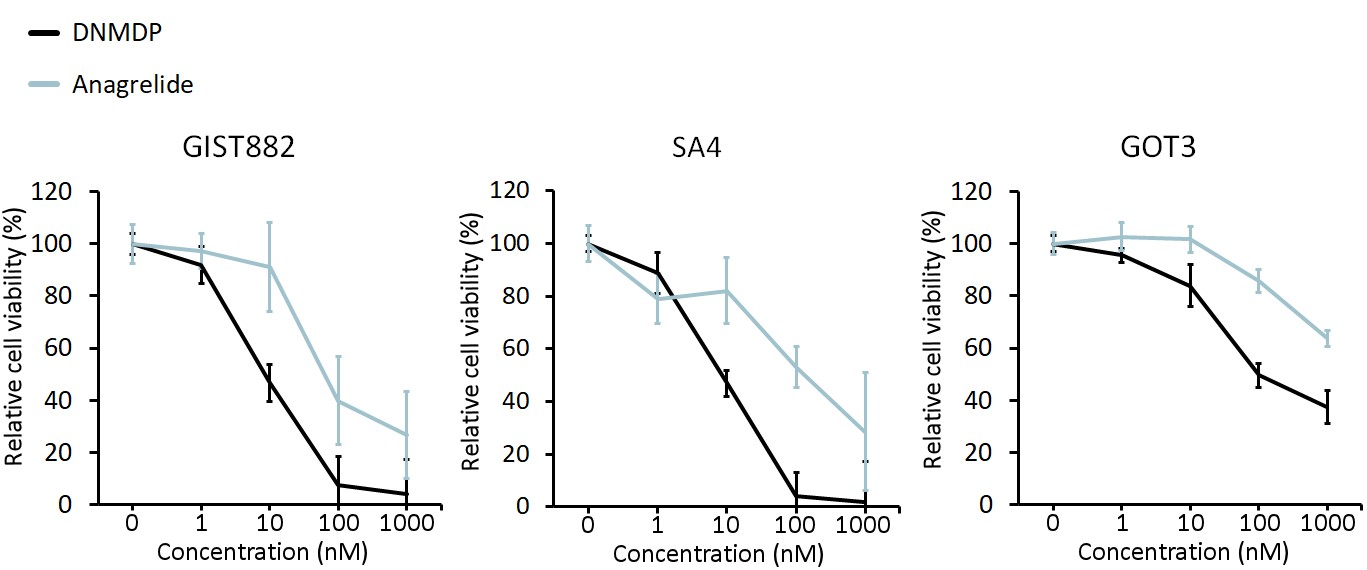

**Figure S4**. Dose-response curves of PDE3A modulators, DNMDP and ANA, in GIST882, SA4, and GOT3 cell lines.

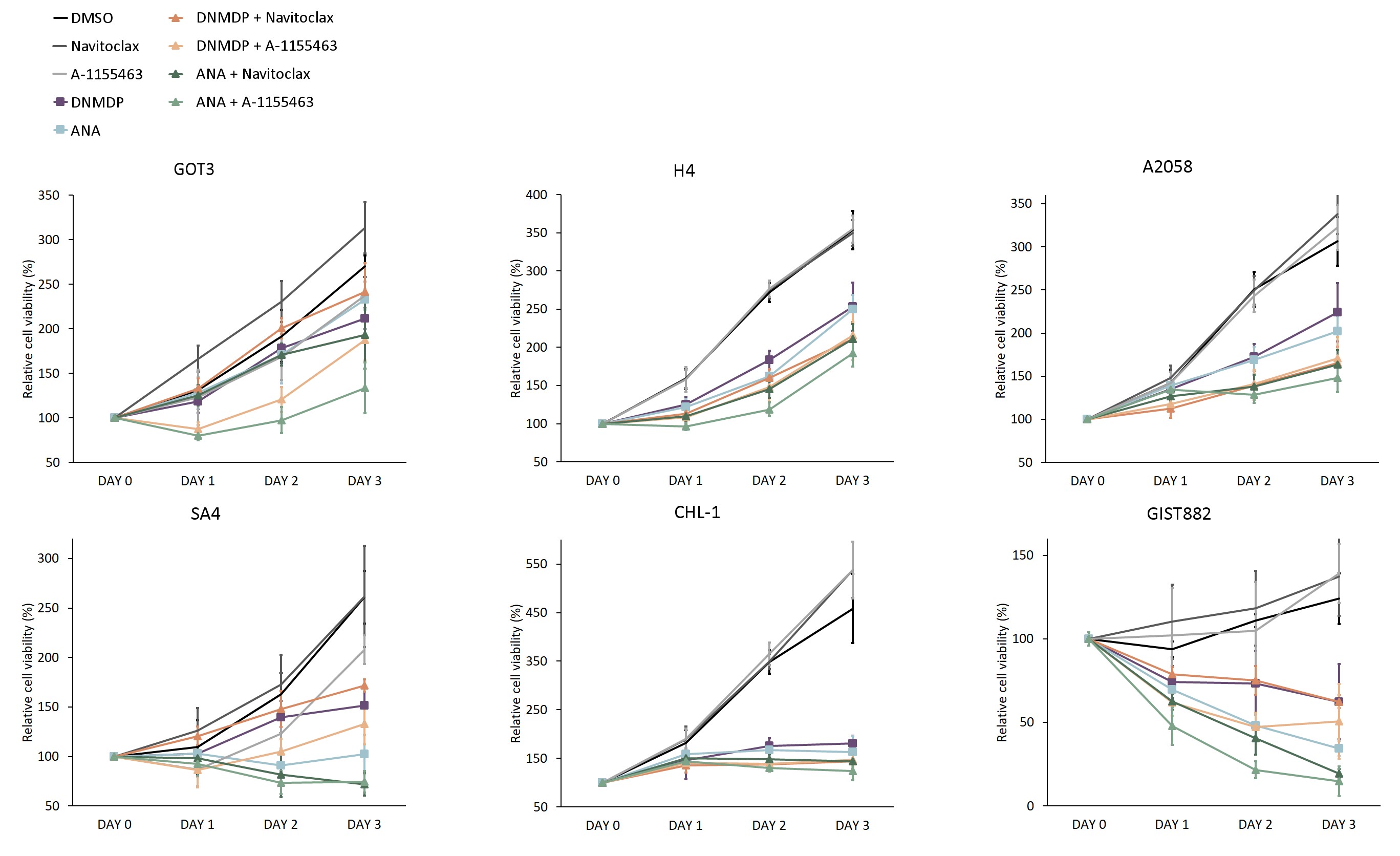

**Figure S5.** Relative cell viability changes through the 72-hour treatment combinations of two PDE3A modulators, ANA and DNMDP, and two Bcl-2/-xL inhibitors, Navitoclax and A-1155463. The combination treatments resulted in the lowest cell viabilities across all cell lines. Abbreviations: ANA = anagrelide.

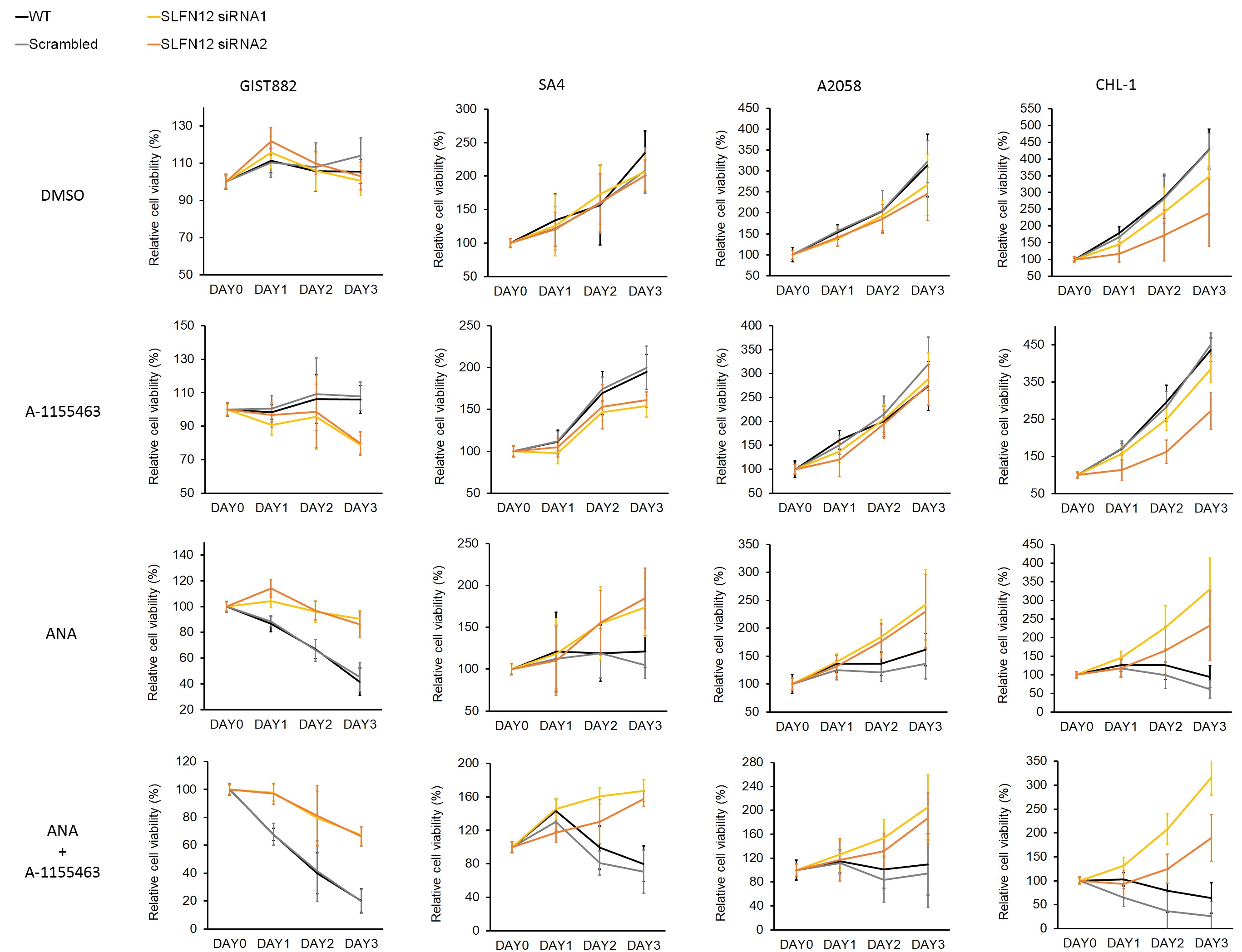

**Figure S6.** SLFN12 silencing using siRNA reduced sensitivity towards ANA and the combination treatment in four PDE3A modulator–sensitive cancer cell lines GIST882, SA4, A2058, and CHL-1. Similar effect was not observed with scrambled siRNA sequences. Abbreviations: ANA = anagrelide.

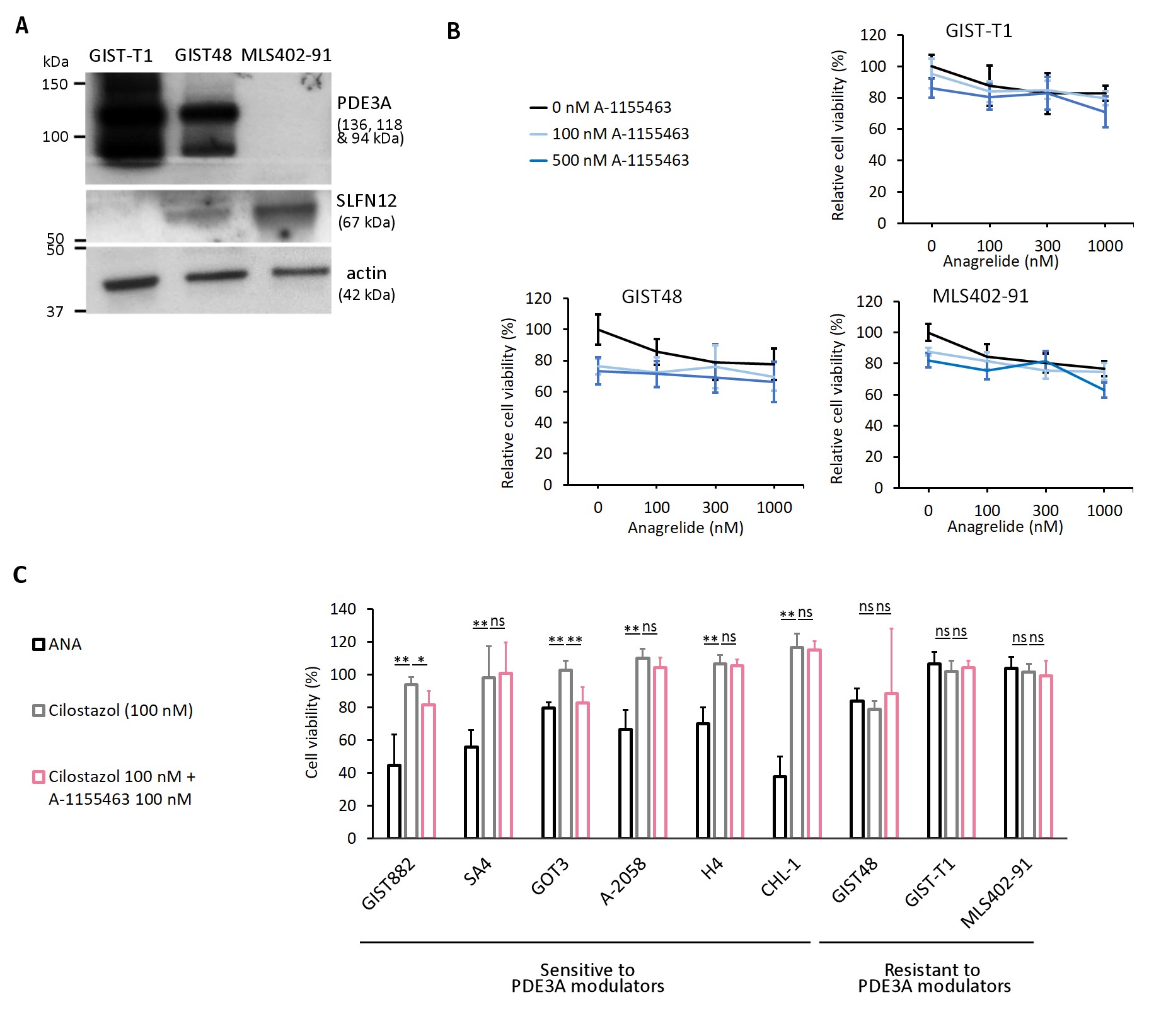

**Figure S7.** **(a)** Immunoblot results of PDE3A and SLFN12 in PDE3A modulator-resistant cell lines; GIST48, GIST-T1, and MLS402-91. **(b)** Dose-response curves of A-1155463 and ANA in the cell lines. Data are shown as mean ± standard deviation from two independent experiments, analyzed using one-way ANOVA with Tukey’s multiple comparison test: *p < 0.01, **p < 0.001, ns = not significant. Abbreviations: ANA = anagrelide.

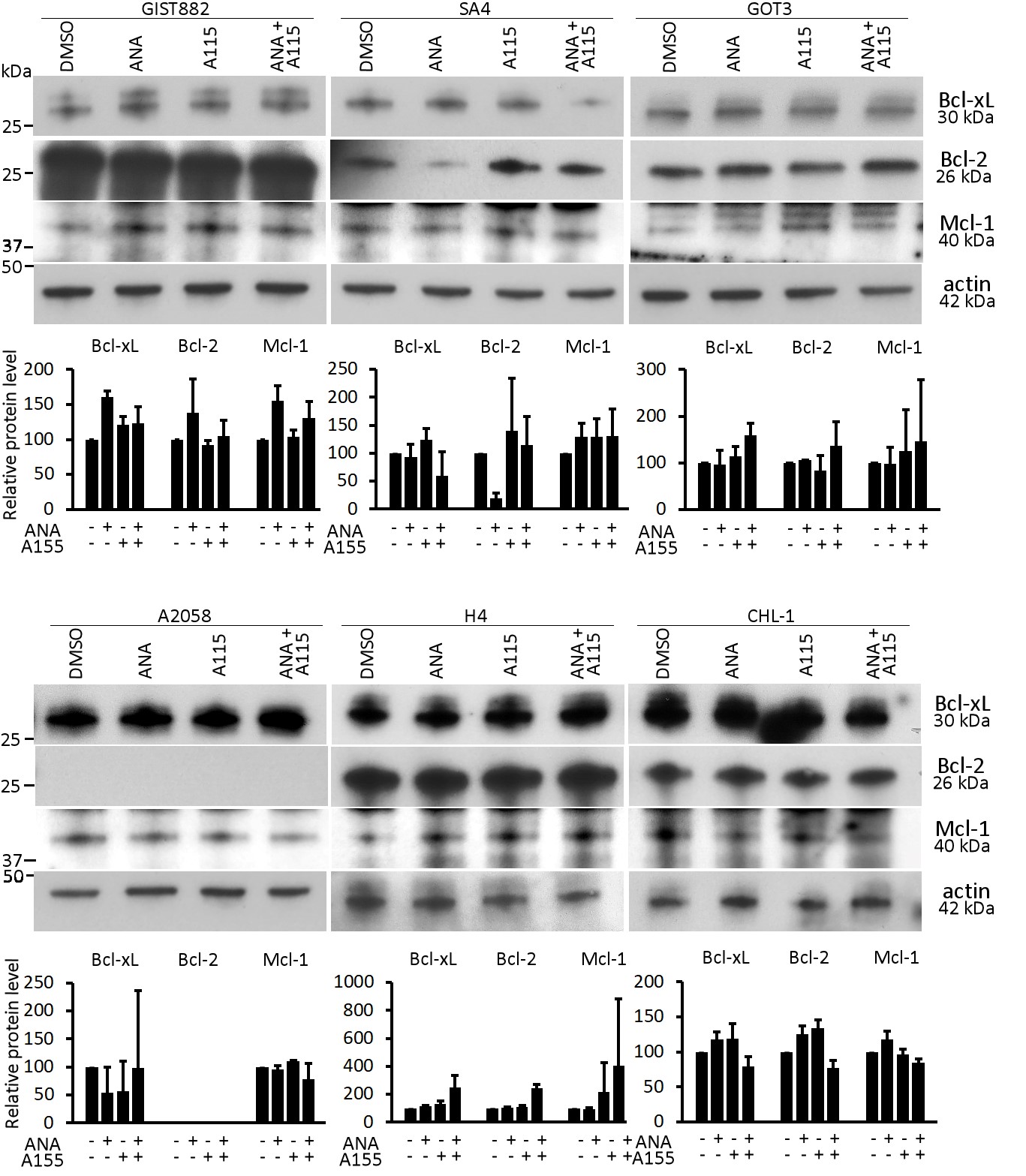

**Figure S8.** Immunoblot and densitometry results for Bcl-xL, Bcl-2, and Mcl-1 were obtained in PDE3A modulator-sensitive cell lines treated with ANA, A-1155463, or both. The treatments did not uniformly affect the expression levels of Bcl-2 family proteins across all cell lines. Abbreviations: ANA = anagrelide; A115 = A-1155463.

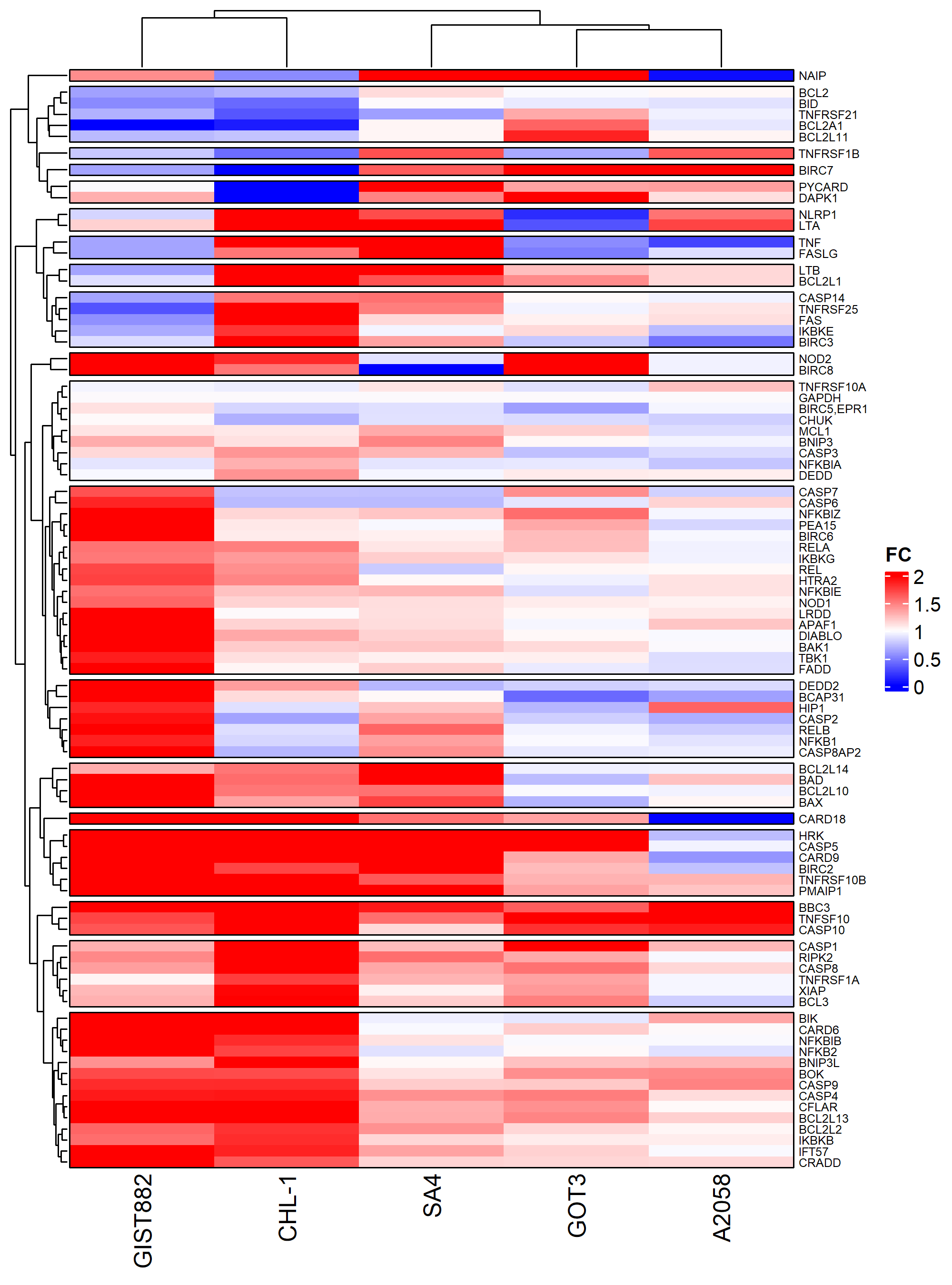

**Figure S9.** A cluster heat map presenting gene expression fold-change values of 89 apoptosis pathway genes after a 24-hour ANA treatment in GIST882, CHL-1, SA2, GOT3, and A2058 cells. To reduce the effect of outliers, the values were capped at a maximum threshold of 2. Abbreviations: FC = fold change.

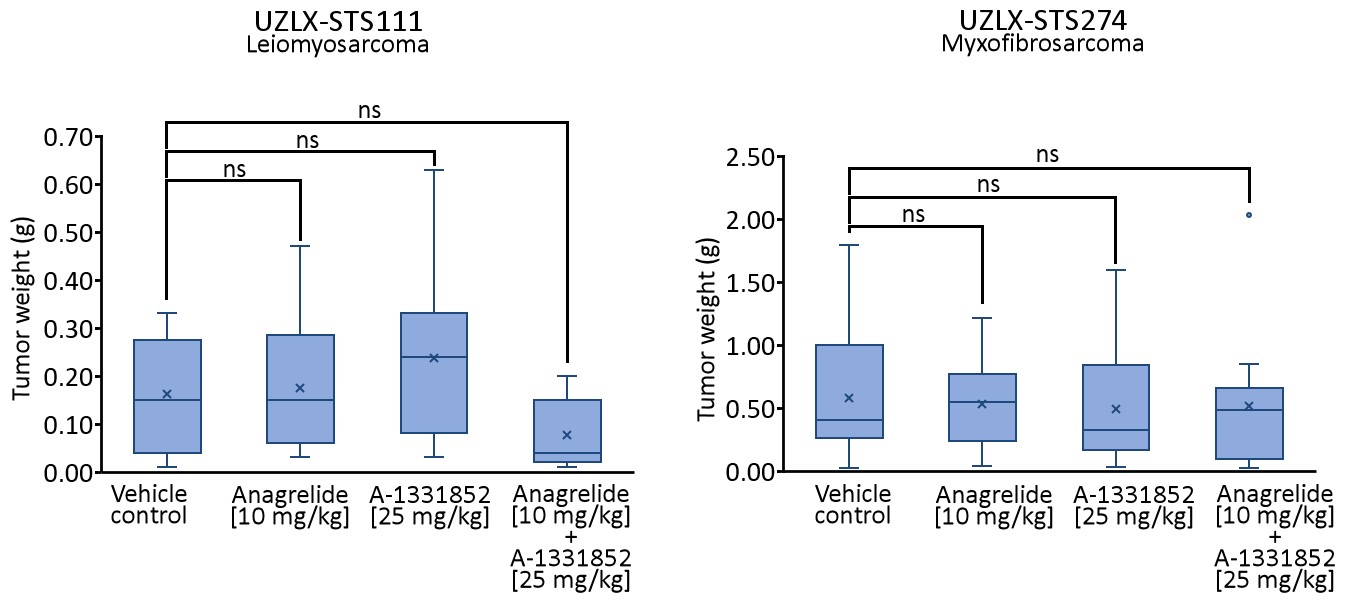

**Figure S10.** Tumor weights at time of collection at the end of the experiment in sarcoma PDX mouse models UZLX-STS111 and UZLX-STS274 treated with ANA, A-1133852, or both. While the combination-treated UZLX-STS111 model tumors weighed on average the least at collection, no significant differences (ns) were detected when analyzed using the Independent-samples Kruskal–Wallis test in either of the models.

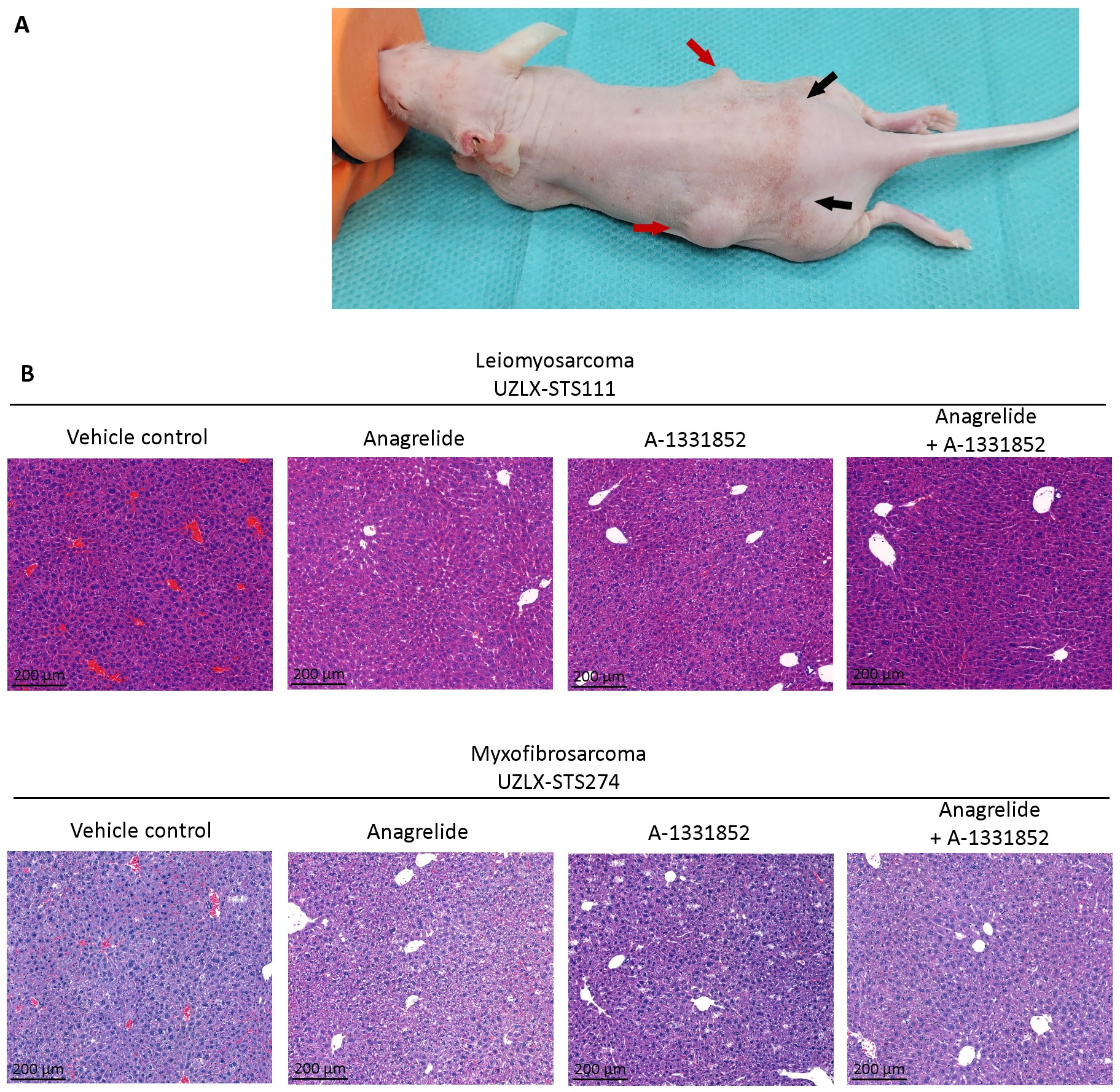

**Figure S11. (a**) All mice in both experiments that received A-1133852 were observed to have red spots on their skin (black arrow). Bilaterally implanted tumors are shown with red arrows. (**b**) Hepatotoxicity was not observed from H&E-stained mouse livers.

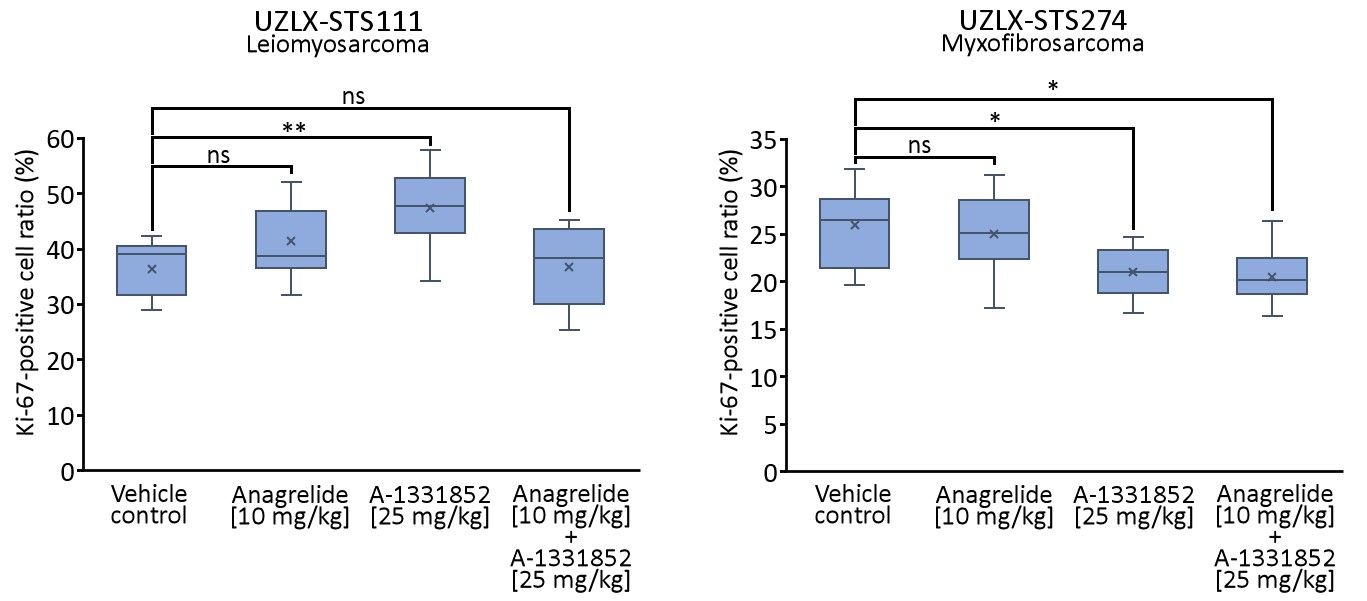

**Figure S12.** Ki-67 positive cell ratios at time of collection at the end of the experiment in sarcoma PDX mouse models UZLX-STS111 and UZLX-STS274 treated with ANA, A-1133852, or both. Although tumor regression was significant in the combination-treated UZLX-STS111 tumors, significant reduction in Ki-67-positive cell ratios, analyzed using one-way ANOVA and Tukey’s multiple comparison test, were not detected. *p < 0.01; **p < 0.001.

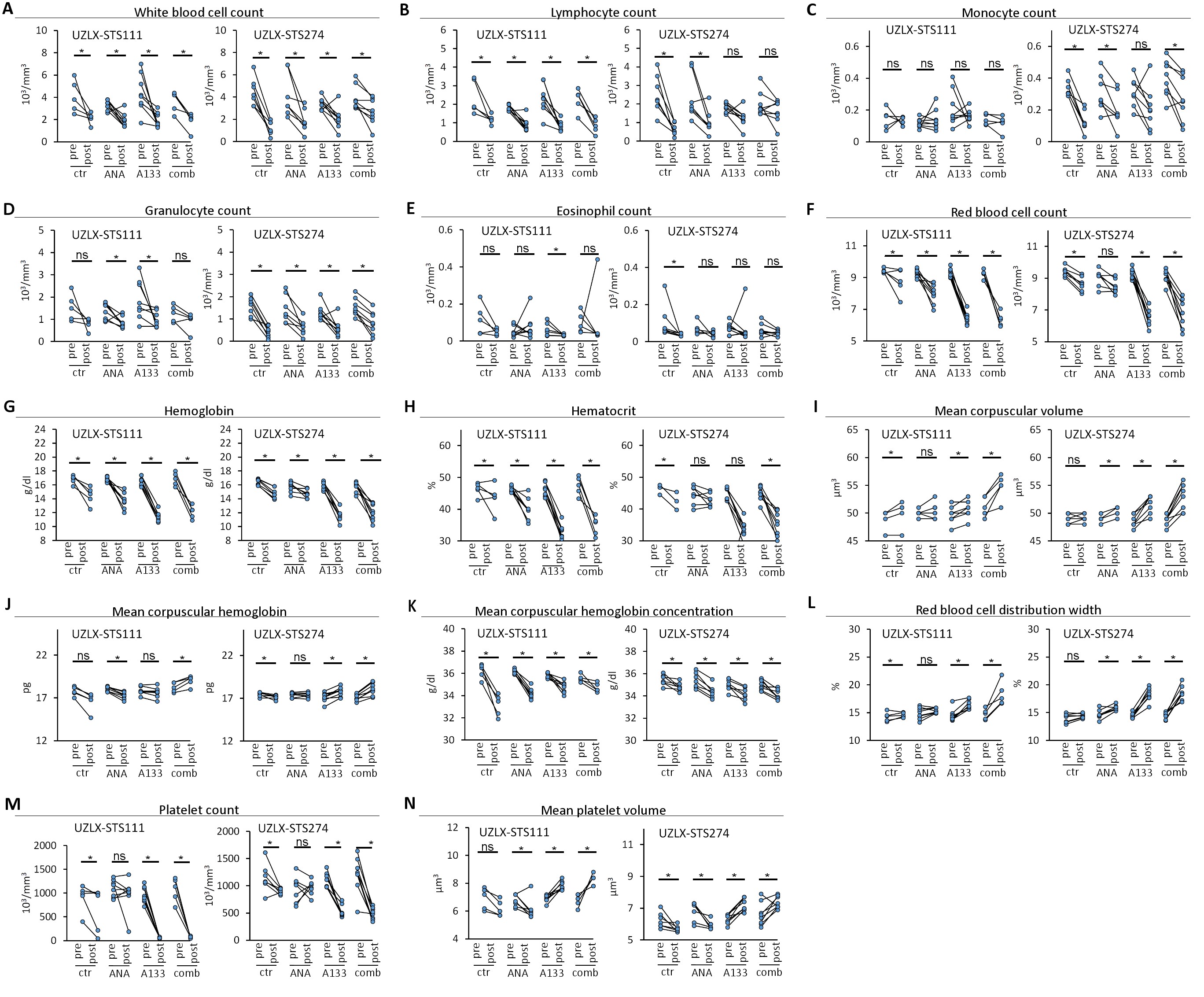

**Figure S13.** Hematological parameters before (pre) and after (post) *in vivo* experiments. Each dot and line represent the change in (**a**) white blood cell, (**b**) lymphocyte, (**c**) monocyte, (**d**) granulocyte, (**e**) eosinophil, (**f**) red blood cell, (**g**) hemoglobin, (**h**) hematocrit, (**i**) mean corpuscular volume, (**j**) mean corpuscular hemoglobin, (**k**) mean corpuscular hemoglobin concentration, (**l**) red blood cell distribution width, (**m**) platelet, and (**n**) mean platelet volume levels in one mouse. Significant changes in blood parameters were analyzed using paired-samples Wilcoxon signed-rank test. Abbreviations: ANA = anagrelide; A133 = A-1331852. *p < 0.05; ns = not significant.
